# Hot-wiring clathrin-mediated endocytosis in human cells

**DOI:** 10.1101/061986

**Authors:** Laura A. Wood, Nicholas I. Clarke, Sourav Sarkar, Stephen J. Royle

## Abstract

Clathrin-mediated endocytosis (CME) is the major route of receptor internalization at the plasma membrane. Analysis of constitutive CME is complicated by the fact that initiation of endocytic events is unpredictable. When and where a clathrin-coated pit will form and what cargo it will contain are difficult to foresee. Here we describe a series of genetically encoded reporters that allow the initiation of CME on demand. This is achieved by inducibly tethering a clathrin “hook” to a plasma membrane “anchor”. Our design incorporates temporal and spatial control of initiation using chemical and optical tools, and the cargo is defined. Since this system bypasses multiple steps in vesicle creation, we term it “hot-wiring”. In this paper, we use hot-wired endocytosis to define the functional interactions between clathrin and the β2 subunit of the AP2 complex. However, there are numerous applications for this new technology, which we hope will be broadly useful to the field.

## Introduction

Clathrin-mediated endocytosis (CME) is the major uptake pathway in eukaryotic cells, which influences numerous processes from nutrition and signaling, to organelle biogenesis and cell excitability (Kirchhausen et al., 2014; Robinson, 2015). In clathrin-mediated synaptic vesicle retrieval, the endocytosis of defined cargo is coupled temporally and spatially to an exocytic event (Granseth et al., 2006; Rizzoli, 2014). By contrast, in constitutive CME, the location of clathrin-coated pit formation in space and time is unpredictable (Ehrlich et al., 2004). Moreover, the cargo contained in each vesicle and the proteins contributing to the inner layer of the clathrin coat are variable (Borner et al., 2012; Taylor et al., 2011). This means that we do not know for certain when or where a vesicle will form, nor what cargo it will contain. Our goal therefore was to design a synthetic system that can be used to trigger endocytosis “on demand”. The aim was to provide temporal and spatial control over the initiation of endocytosis, using defined cargo.

A straightforward method to trigger endocytosis is the activation of a receptor at the cell surface, for example GPCRs activated by their cognate ligand (Puthenveedu et al., 2007). This would provide temporal control and could be adapted for spatial control; however, i) activation of intracellular signaling would complicate analysis, ii) the precise molecular details for activation-dependent internalization of many receptors are not completely understood, iii) this would not report on constitutive CME, and iv) activated GPCRs may not generate clathrin-coated pits *de novo* (Lampe et al., 2014). For this reason, we sought a synthetic system to initiate CME.

The major clathrin adaptor at the plasma membrane is the AP2 complex. AP2 performs the essential function of recognizing cargo and membrane, and also contacts clathrin via its β2 subunit, specifically the hinge and appendage domains (Keyel et al., 2008; Murphy and Keen, 1992). The AP2 complex undergoes a number of large-scale conformational changes in order to bind membrane, recognize cargo and become ready for clathrin engagement (Jackson et al., 2010; Kelly et al., 2014; Kelly et al., 2008). In designing a synthetic system to trigger endocytosis on demand, these regulatory steps would need to be bypassed so that the process can be “hot-wired”.

Elegant *in vitro* studies show that clathrin-coated pits can be formed by anchoring a clathrin binding protein (a clathrin “hook”) at a membrane (Dannhauser and Ungewickell, 2012). We reasoned that a similar approach, if it could be made to be inducible, would trigger endocytosis inside living, human cells. This paper describes our design and optimization of synthetic reporters to trigger endocytosis on demand in human cells. We show that this system can be applied to answer specific cell biological questions, such as defining the molecular requirements for clathrin-AP2 interaction.

## Results

### Synthetic, constitutively active endocytosis as a proof-of-principle

As a first step towards hot-wiring endocytosis, we created a fusion protein to serve as a proof-of-principle. This fusion, of the appendage and hinge of β2 with a single-pass transmembrane protein (CD8-β2-mCherry, Figure 1A), was effectively a clathrin hook anchored at the plasma membrane, and we tested its ability to be internalized. Using live cell antibody feeding we observed endocytosis of this construct at physiological temperatures, but not at 4 °C (Figure 1B, C). The single-pass transmembrane protein (CD8α) in the absence of the fusion cannot be internalized (Fielding et al., 2012; Kozik et al., 2010). These proof-of-principle experiments confirmed that the presence of a clathrin hook at the plasma membrane was sufficient for a transmembrane protein to be internalized.

**Figure 1.**
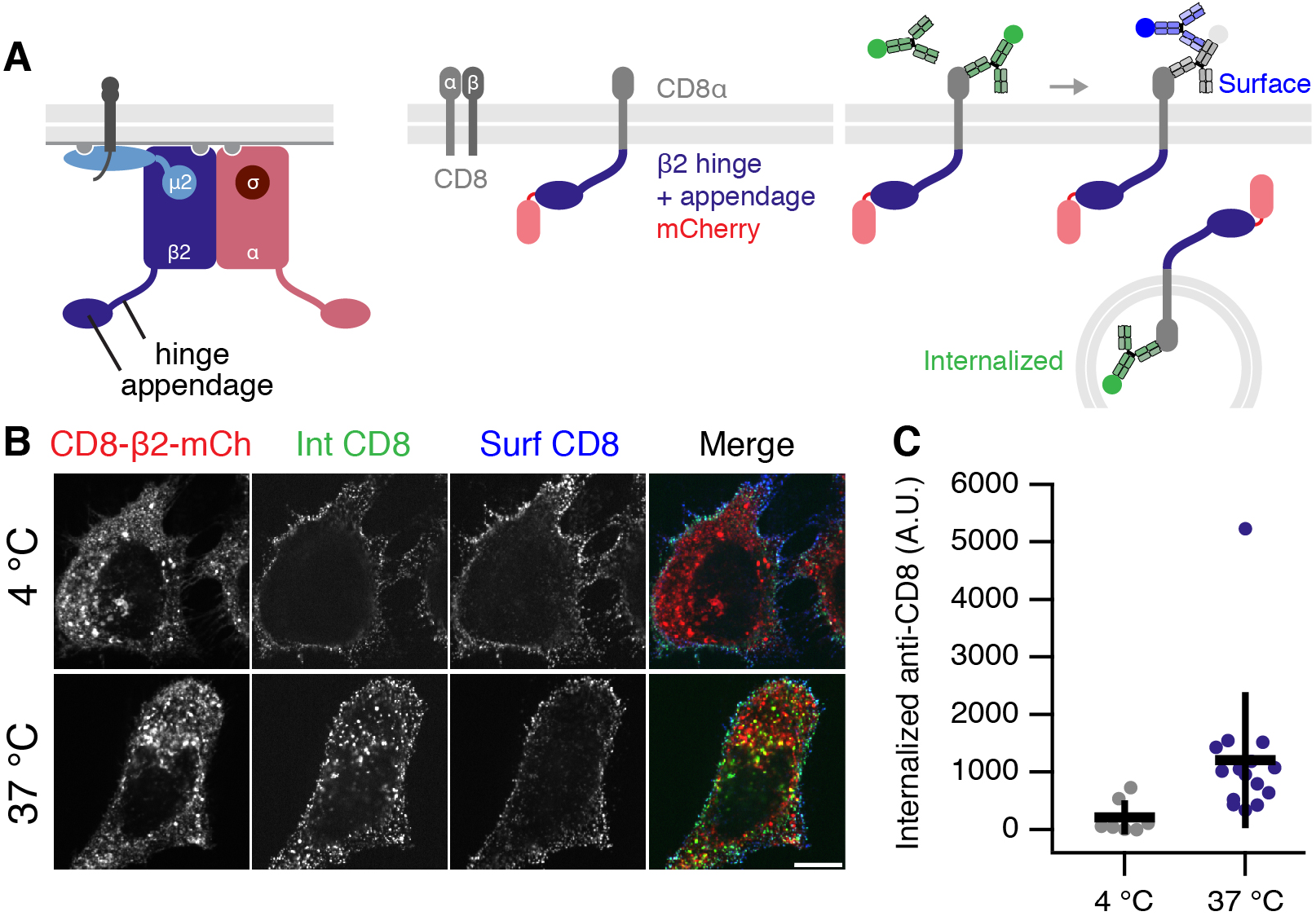
Proof-of-principle that clathrin-mediated endocytosis can be hot-wired. (A) Schematic diagram of AP-2 complex comprising α, β2, µ2 and σ2 subunits. Note that β2 has a hinge and appendage which must be uncovered for clathrin engagement. Center, CD8 is a heterodimer of α and β subunits. In CD8-β2-mCherry, the CD8a chain is directly fused to the β2 hinge and appendage and tagged with mCherry. Right, antibody feeding assay. Internalized CD8-β2-mCherry is detected by Alexa488-conjugated anti-CD8 (see Methods). (B) Representative confocal micrographs of live immunolabeling experiment to demonstrate constitutive internalization of CD8-β2-mCherry. Cells were labeled with Alexa488-conjugated anti-CD8 and then incubated at 4 °C or 37 °C for 40 min. Surface signal is quenched and re-stained with Alexa633-conjugated secondary antibody. Scale bar, 10 μm. (C) Quantification of internalized CD8-β2-mCherry at 4 °C and 37 °C. Student's t-test, p = 0.0067. Bars show mean ± 1 s.d.

### Development of chemically inducible endocytosis

We next designed a series of constructs that would allow us to hot-wire endocytosis (Figure 2). The FKBP-rapamycin-FRB system was exploited to induce the dimerization of a plasma membrane anchor with a clathrin hook and thereby control the initiation of endocytosis (Figure 2A). Live cell imaging demonstrated that the clathrin hook was rapidly recruited to the plasma membrane in response to rapamycin. Immediately afterwards, bright green puncta began to form. These bright puncta only occurred when a clathrin hook (FKBP-β2-GFP) was rerouted to the plasma membrane, but not when a construct lacking the hook (GFP-FKBP) was used (Figure 2B, C, Supplementary Movies S1 and S2). Similar responses were detected using a plasma membrane anchor based on the monomeric transmembrane protein CD4 and even with a palmitoylated peripheral membrane protein acting as the anchor (GAP43-FRB-mRFP) (Figure 2-figure supplement 1). Interestingly, no bright green puncta were formed when a clathrin hook was sent to the mitochondria indicating that puncta formation depends on the plasma membrane and on anchors that are correctly addressed there (Figure 2-figure supplement 2).

**Figure 2.**
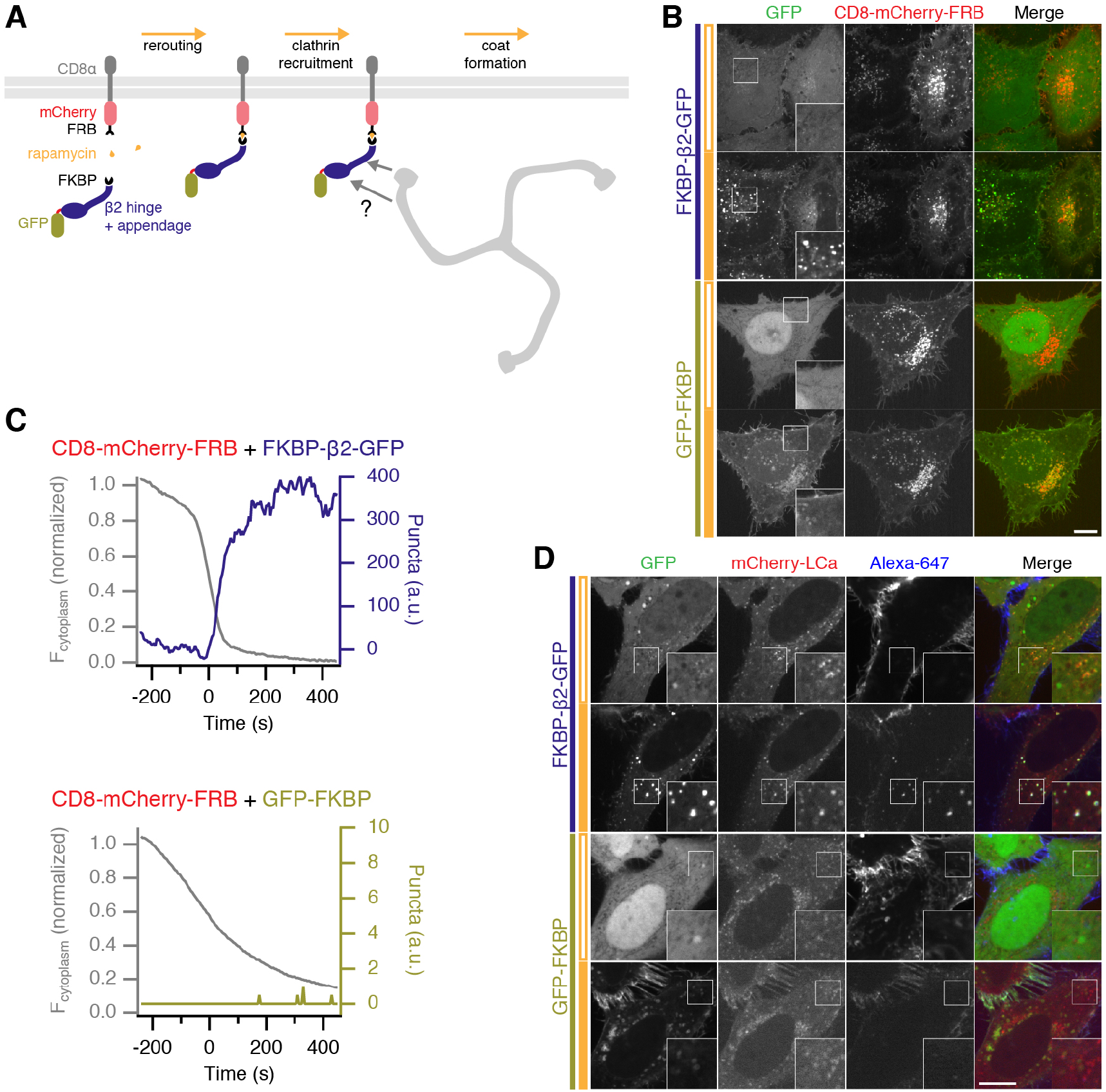
Chemically-inducible endocytosis. (A) Illustration of chemically-inducible endocytosis. Cells co-express a plasma membrane anchor (CD8-mCherry-FRB) and a clathrin hook (FKBP-Β2-GFP). Rapamycin (200 nM) is added which causes heterodimerization of FKBP and FRB domains. The clathrin hook is rerouted to the plasma membrane. Clathrin recognizes the clathrin hook and the plasma membrane anchor is internalized. (B) Chemically-inducible endocytosis in live cells. Cells expressing CD8-mCherry-FRB with either FKBP-β2-GFP or GFP-FKBP. Stills from live cell imaging experiments are shown, upper image shows the frame before rerouting occurs and lower image is 133 frames (665 s) later. See Supplementary Movies S1 and S2. (C) Example plots to show chemically-inducible endocytosis. Colored traces indicate the total area of bright GFP puncta that form in cells after rerouting. Gray traces show the rerouting of the clathrin hook from the cytoplasm to the plasma membrane (see Methods).
(D) Bright green spots formed by chemically-induced endocytosis colocalize with clathrin and contain anti-CD8-Alexa647. Cells co-expressing mCherry-LCa, CD8-dCherry-FRB and either FKBP-β2-GFP or GFP-FKBP, were incubated with anti-CD8-Alexa647 to label the membrane anchor extracellularly, and rapamycin (200 nM) was applied. In B and D, insets show a 2X zoom of the boxed region; closed orange bar indicates rapamycin application; scale bars, 10 μm.

**Figure 2-figure supplement 1.**
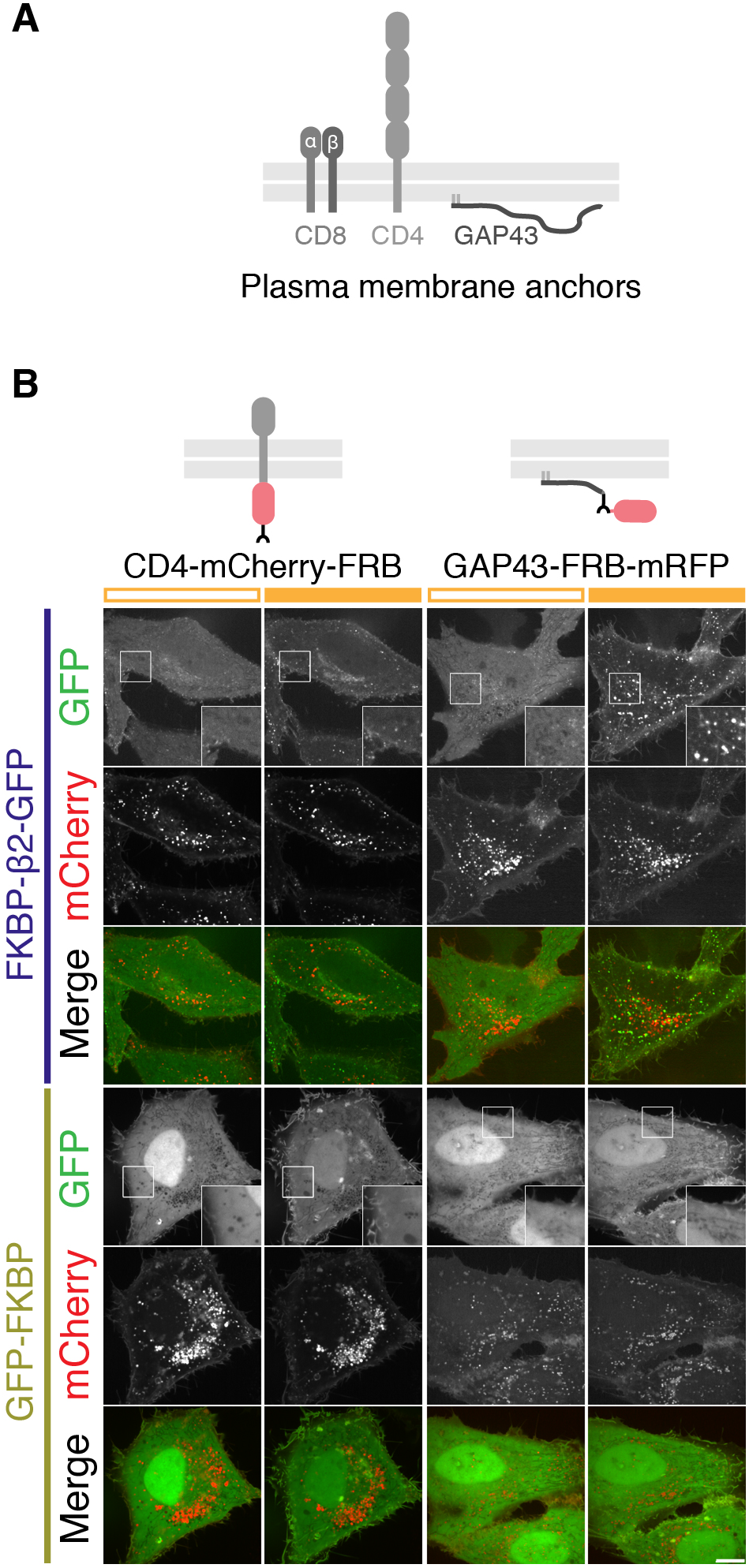
Plasma membrane anchors are interchangeable for chemically-induced internalization. (A) Schematic layout of CD8, CD4 and GAP43, used for engineering plasma membrane anchors. (B) Comparison of two other plasma membrane anchors used for chemically-inducible endocytosis. Cells expressed CD4-mCherry-FRB or GAP43-FRB-mRFP with either FKBP-β2-GFP or GFP-FKBP. Stills from live cell imaging experiments are shown, left image shows the frame before rerouting occurs and right image is 133 frames (665 s) later. Both anchors behaved similarly to CD8-mCherry-FRB. Inset shows a 2X zoom of the boxed region. Scale bar, 10 μm.

**Figure 2-figure supplement 2.**
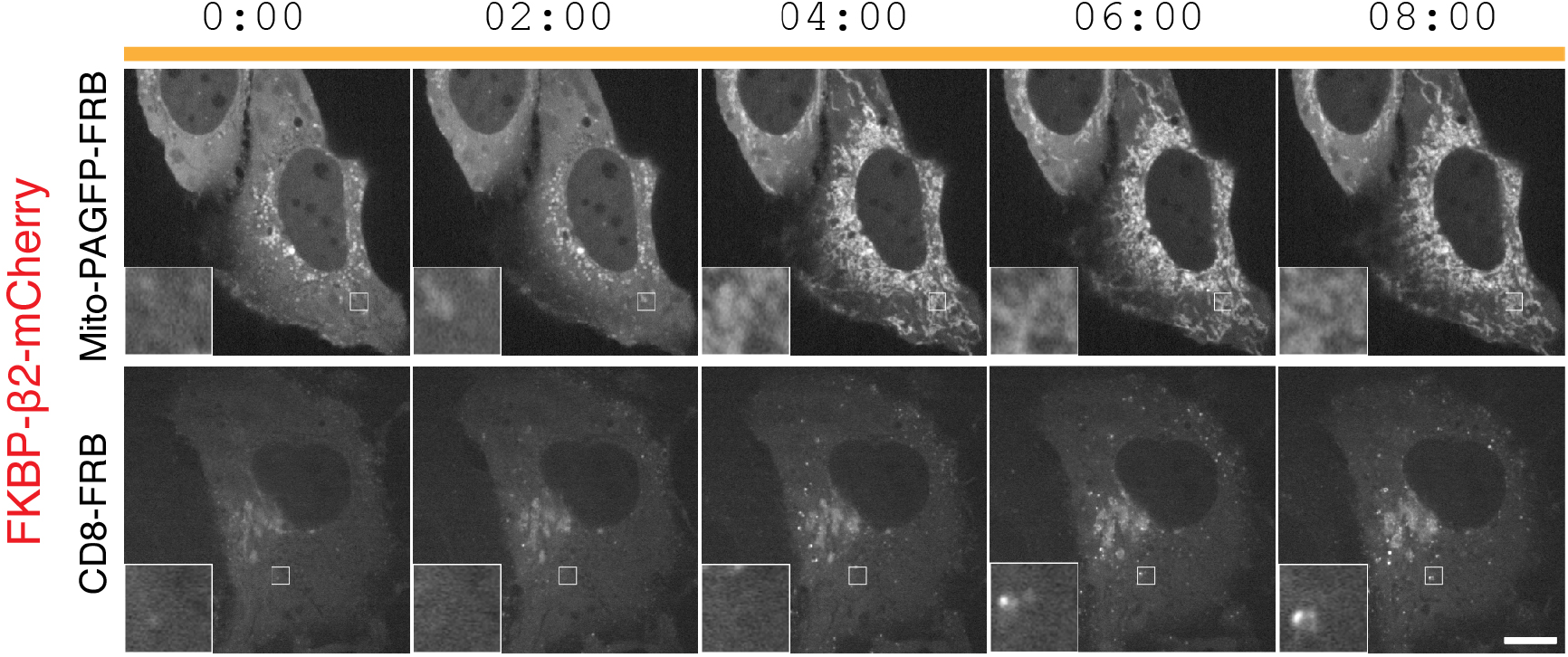
Sending a clathrin hook to a mitochondrial anchor does not result in vesicle formation. Stills from a typical live cell confocal imaging experiment. HeLa cells expressing a mitochondrial anchor (Mito-PAGFP-FRB) or a plasma membrane anchor (CD8-FRB) were compared using a clathrin hook (FKBP-β;2-mCherry). Rapamycin (200 nM) was added at time zero. Insets show 5X zoom of the boxed area. Scale bar, 10 μm.

**Figure 2-figure supplement 3.**
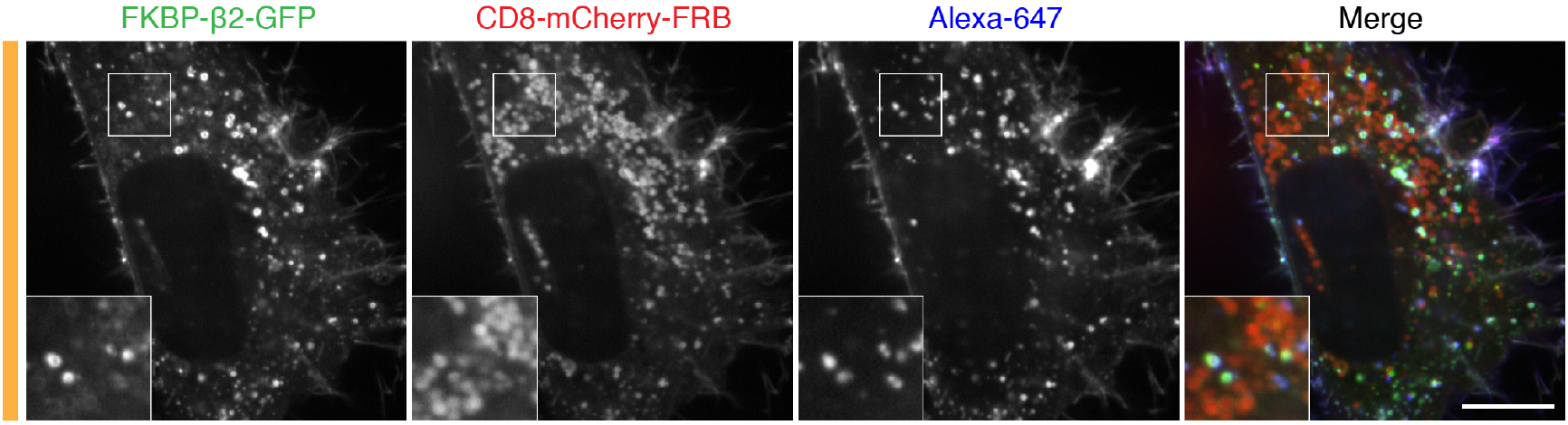
Bright green puncta originate from the plasma membrane and can be distinguished from other spots. Still from a typical live cell confocal imaging experiment. HeLa cells co-expressing CD8-mCherry-FRB and FKBP-β2-GFP were incubated with anti-CD8-Alexa647 to label the membrane anchor extracellularly, and rapamycin (200 nM) was applied. Insets show 2X zoom of the boxed area. Scale bar, 10 μm.

Before application of rapamycin, plasma membrane anchors were found in intracellular structures to some extent, in addition to the population at the plasma membrane. The clathrin hook is also initially recruited to these structures forming larger, dimmer spots, but they do not generate bright green puncta (Figure 2B and Supplementary Movie S1). Using antibody feeding, we found that the dimmer structures do not contain extracellularly applied antibody, but that the bright green puncta do (Figure 2-figure supplement 3). Using image analysis methods, it was possible to distinguish the bright green puncta from the large dimmer accumulations at stationary spots present at the start of the experiment (see Methods, Figure 2B). If chemically-induced internalization is indeed a hot-wired form of CME, then these observations indicate that the bright green puncta are a readout of this process.

### Chemically-induced internalization is a hot-wired form of CME

We suspected that the bright green puncta that are generated from the plasma membrane upon heterodimerization of clathrin hook and membrane anchor were clathrin-coated vesicles (CCVs). We tested this hypothesis using four different approaches.

First, we used antibody feeding and simultaneous clathrin imaging in live cells where chemically-induced internalization was triggered. The plasma membrane anchor was labelled extracellularly with anti-CD8/Alexa647 and the heterodimerization of FKBP and FRB was then triggered with rapamycin (Figure 2D). The bright green puncta that form in cells expressing the clathrin hook contained anti-CD8, which confirmed that these puncta are formed by internalization from the plasma membrane. No bright green puncta nor antibody uptake was seen in cells co-expressing GFP-FKBP. Moreover, the bright green puncta also colocalized with clathrin (mCherry-LCa), suggesting that they are clathrin-coated (Figure 2D, Supplementary Movie S3).

**Figure 2-figure supplement 4.**
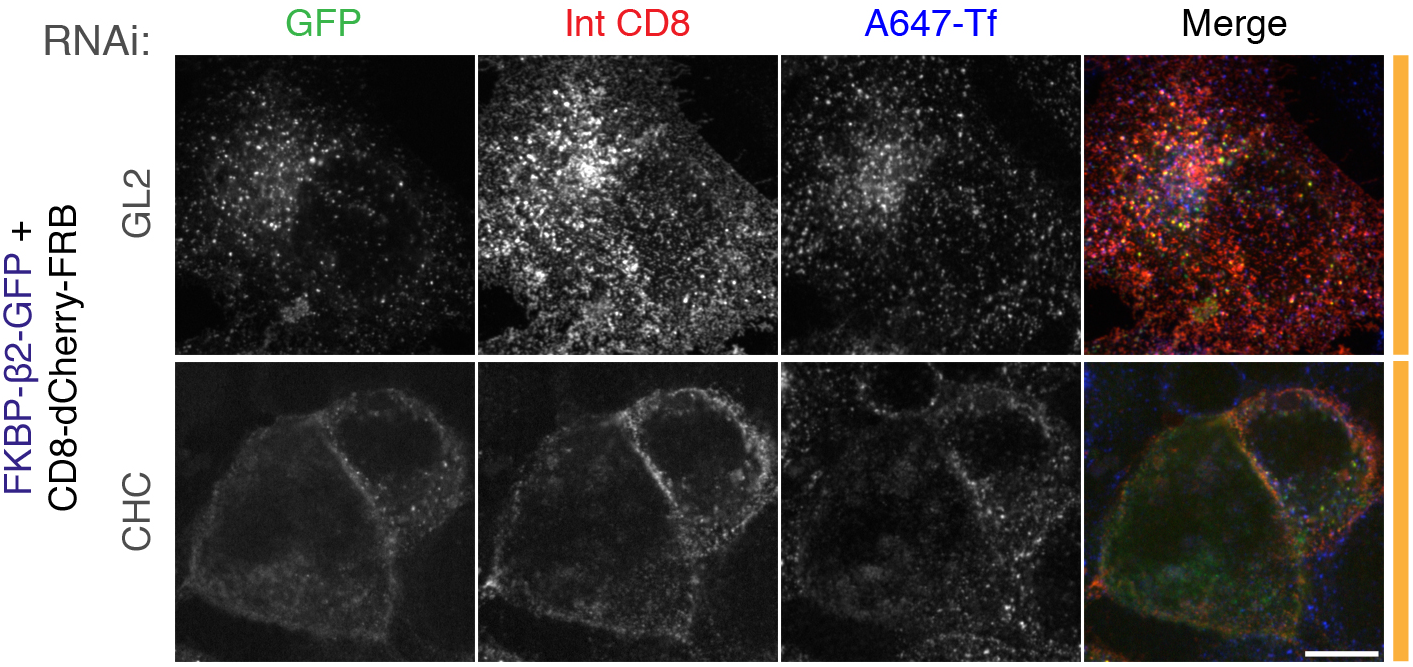
Chemically-induced endocytosis is clathrin-dependent. Live cell immunolabeling experiments of cells expressing CD8-dCherry-FRB (dark mCherry variant) with FKBP-β2-GFP. Cells were also transfected with siRNAs against GL2 (control) or clathrin heavy chain (CHC). Inhibition of uptake of transferrin-Alexa647 (A647-Tf, blue) was used as a functional test of knockdown efficacy. Scale bar, 10 μm.

Second, in cells depleted of clathrin heavy chain using RNAi, rapamycin-induced heterodimerization of anchor and hook did not result in the formation of bright green puncta (Figure 2-figure supplement 4). This result confirms that chemically-induced internalization is clathrin-mediated.

Third, we used total internal fluorescence microscopy (TIRFM) to image chemically-induced internalization at the plasma membrane. Internalization was triggered in RPE1 cells co-expressing CD8-dCherry-FRB, FKBP-β2-mRuby2 and either dynamin2-GFP or GFP-LCa. We observed that the FKBP-β2-mRuby2 puncta that form after rapamycin application, terminate with an accumulation of dynamin consistent with scission (Figure 3). As expected, FKBP-β2-mRuby2 puncta colocalize with clathrin throughout their lifetime from appearance to disappearance (Figure 3). The timecourse of these events was similar to that reported elsewhere (Aguet et al., 2013).

**Figure 3.**
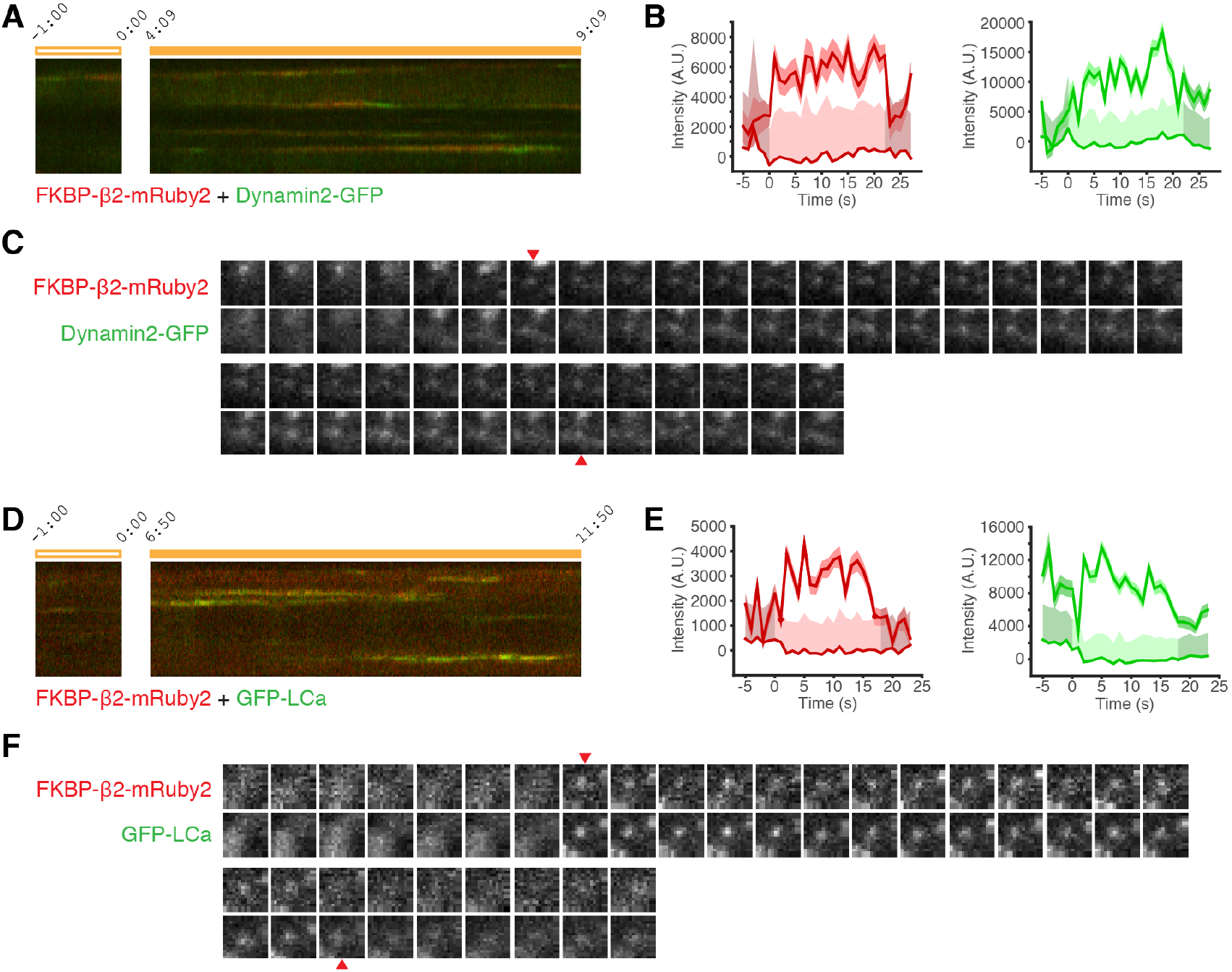
Imaging chemically-inducible endocytosis using TIRFM. Total internal fluorescence microscopy (TIRFM) imaging in RPE1 cells expressing CD8-dCherry-FRB, FKBP-β2-mRuby2 and either dynamin2-GFP (A-C) or GFP-LCa (D-F). Chemically induced endocytosis was triggered by rapamycin (200 nM). (A,D) Kymograph to show the fluorescence along a line in the cell footprint over time. Open and filled orange bars indicate pre and post rapamycin application. (B,E) Two example intensity traces of puncta at the plasma membrane imaged by two-color TIRFM and identified by model-based detection and tracking of FKBP-β2-mRuby2. Plots show for each channel: intensity with a 1 SD uncertainty (shaded area) and background intensity with 2 SD significance threshold (light areas indicate where an event was above threshold, detected). Note the time refers to the event and not to rapamycin application. (C,F) Frame-by-frame montage of the spot analyzed in B and E, respectively. Arrowheads indicate the time when the event was above detection.

Fourth, we tested whether or not the bright puncta were CCVs using correlative light-electron microscopy (CLEM). To do this, we again took advantage of the ability of our system to be tracked using antibodies to the extracellular domain of CD8. HeLa cells co-expressing CD8-dCherry-FRB and either FKBP-β2-GFP or GFP-FKBP as a control, were labeled by anti-CD8/Alexa546-FluoroNanoGold and endocytosis was initiated by addition of rapamycin (200 nM). By fluorescence microscopy, the formation of bright puncta was confirmed in cells co-expressing FKBP-β2-GFP but not GFP-FKBP (Figure 4A). The same cell observed by light microscopy was then fixed, processed and imaged by EM. In cells that formed bright puncta, gold particles were found in CCVs (Figure 4B). These CCVs had a slightly enlarged morphology compared with regular CCVs (diameter of 114.7 ± 3.8 nm *vs* 91.0 ± 2.2 nm, mean ± s.e.m., p = 2.1 × 10^−6^, Figure 4C). This difference was mainly due to the thickness or density of the coat surrounding the vesicle (46.1 ± 2.0 *vs* 31.2 ± 1.1 nm, p = 4.7 × 10^−8^). In controls, gold particles remained at the cell surface and were not found in CCVs (Figure 4B).

**Figure 4.**
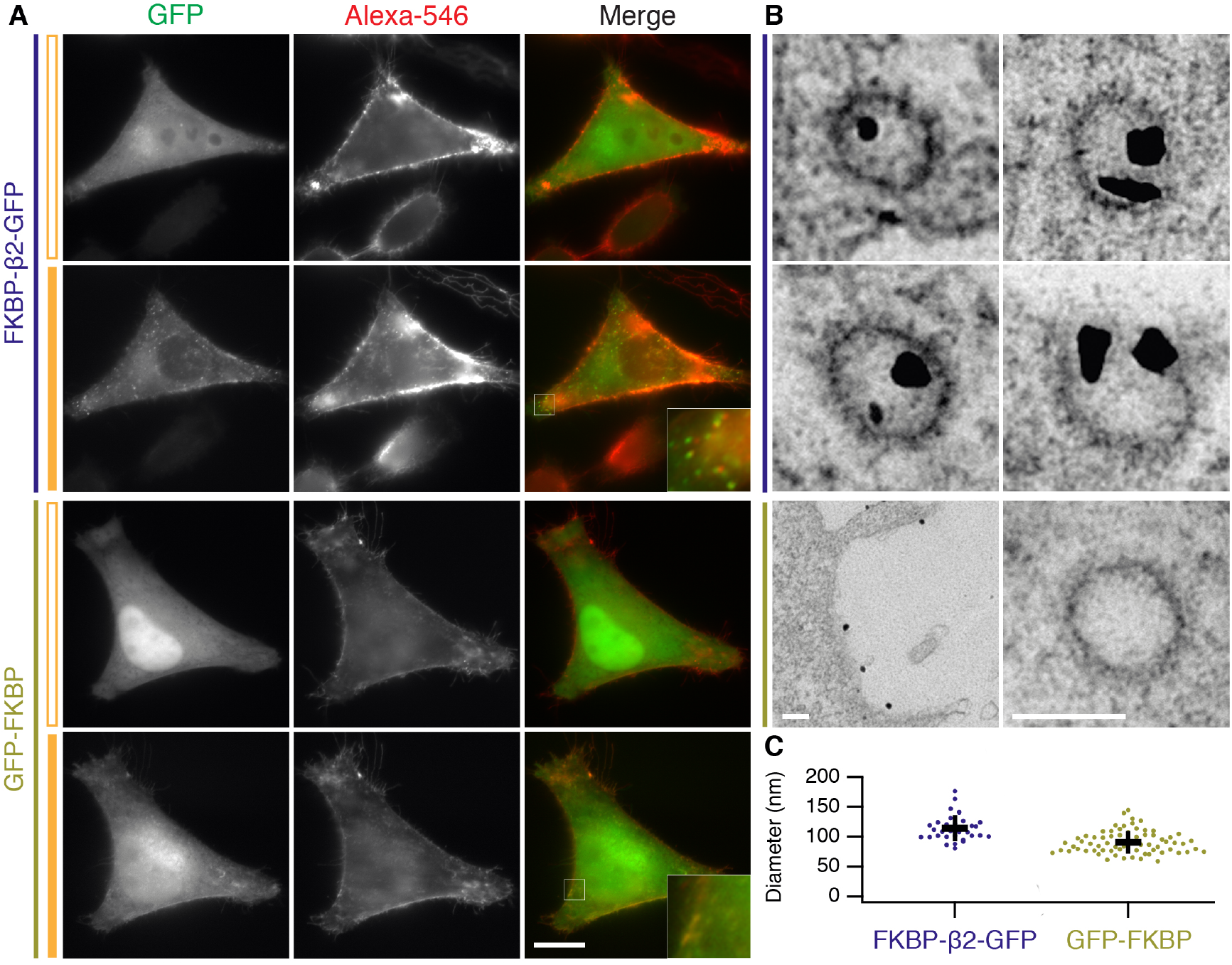
Chemically-induced internalization is via clathrin-coated vesicles. Correlative light-electron microscopy to study the uptake of immunolabeled CD8-dCherry-FRB in cells co-expressing either FKBP**-β**2-GFP or GFP-FKBP. Results from a single experiment are shown, although CLEM experiments were performed three times. (A) Stills from live cell imaging showing cells sequentially labeled with anti-CD8 and Alexa546 FluoroNanoGold conjugated secondary antibody, before and after addition of rapamycin (200 nM). The same cell was processed for EM and imaged. Scale bars, 10 μm. (B) Electron micrographs of the cells shown in A. Clear uptake of NanoGold into CCVs was seen in cells co-expressing FKBP**-β**2-GFP but not GFP-FKBP. Scale bars, 100 nm. (C) Scatter plot of CCV diameters in FKBP**-β**2-GFP versus GFP-FKBP samples. Note CCVs in FKBP**-β**2-GFP contained NanoGold, whereas GFP-FKBP CCVs did not. Student's t-test, p ± 2.1 × 10^−6^. Bars indicate the mean ± 1 s.d.

This result confirms that chemically-induced internalization is via a clathrin-dependent mechanism.

Together, these experiments show that chemically-induced internalization is a hot-wired form of CME because i) internalization occurs at the plasma membrane and ii) is clathrin-dependent, iii) the vesicles are clathrin-coated and iv) their scission is concomitant with dynamin recruitment. Therefore, the bright green puncta which form upon chemically-induced endocytosis are a readout of hot-wired CME.

### Is hot-wired endocytosis orthogonal to constitutive CME?

We next wanted to test if hot-wired endocytosis is orthogonal to normal CME. That is, does the introduction and/or application of such a synthetic system interfere with constitutive membrane traffic? To address this, we monitored transferrin uptake in cells expressing FKBP**-β**2-GFP alone or with CD8-mCherry-FRB. Transferrin uptake appeared normal in the presence or in the absence of rapamycin (Figure 5A, B). This indicates that triggering synthetic endocytosis is not sufficient to block regular CME. Moreover, there was no obvious detrimental effect on CME by merely expressing our system in cells (Figure 5A, B). In a second test of orthogonality, we found that triggered endocytosis occurred in cells that were depleted of the μ2 subunit of AP2 (Figure 5C, D). The depletion by RNAi was good (Figure 5E) and was sufficient to block endogenous transferrin uptake. These experiments suggest that hot-wired endocytosis can operate orthogonally, that is, independent of and without gross detriment to, endogenous CME. However, the extent of orthogonality and how hot-wired events interface with regular CME in normal cells may need further investigation.

**Figure 5.**
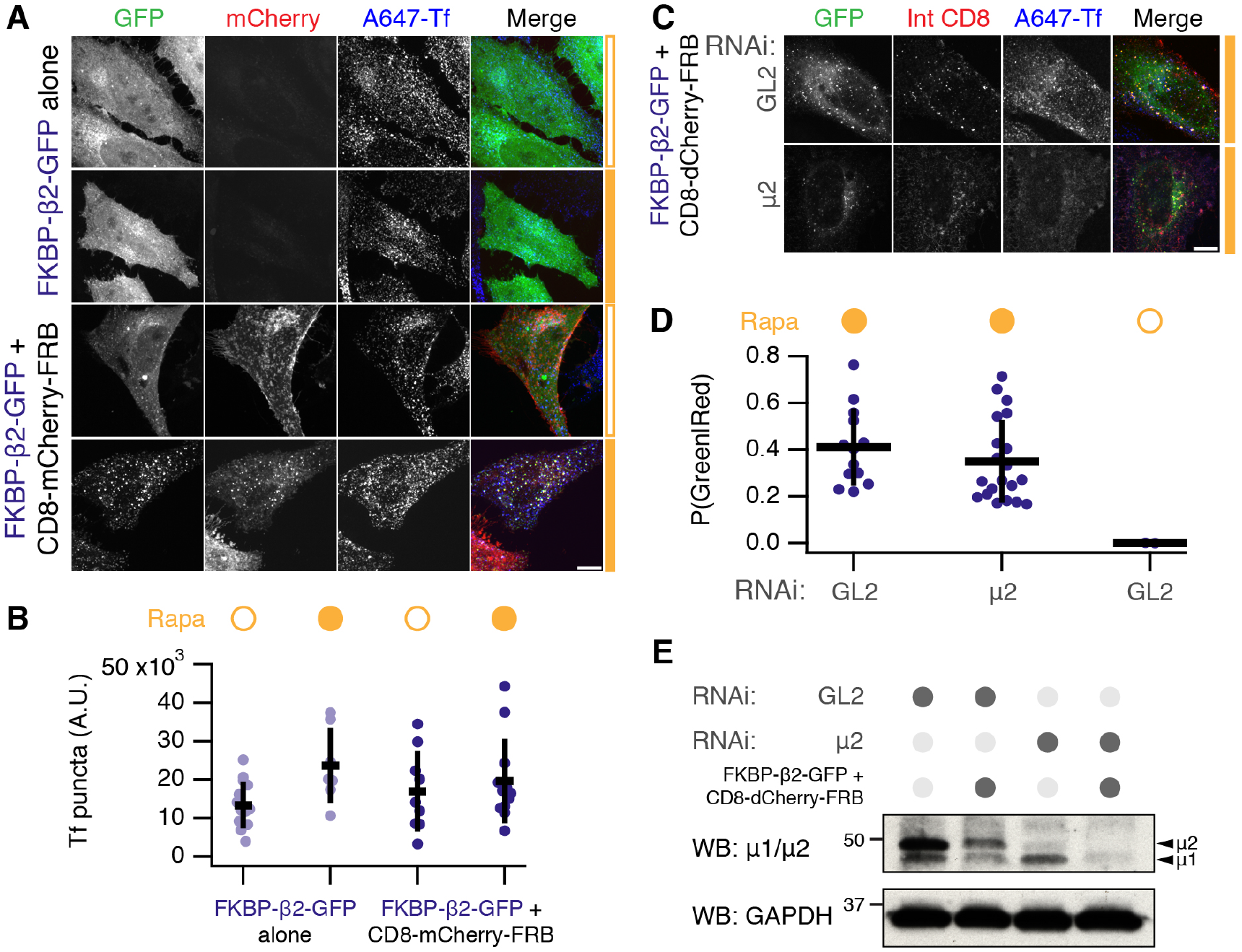
Hot-wired endocytosis can operate orthogonally to normal CME. (A) Orthogonality of chemically-inducible endocytosis. HeLa cells expressing FKBP-β2-GFP alone or together with CD8-mCherry-FRB were tested for ability to internalize transferrin-Alexa647 (A647-Tf) in the presence or absence of rapamycin (200 nM, filled orange bar). (B) Quantification of transferrin uptake and number of GFP-positive puncta in cells. Scatter plot shows values for each cell, bars indicate the mean ± 1 s.d. N_exp_ = 2. Tf, ANOVA, p = 0.13; GFP, ANOVA, p = 5.89 × 10^−6^. (C) Chemically-induced endocytosis can operate independently of endogenous AP-2. Live cell immunolabeling experiments of cells expressing CD8-dCherry-FRB (dark mCherry variant) with FKBP-β2-GFP. Cells were also transfected with siRNAs against GL2 (control) or μ2 subunit of AP-2 complex. Inhibition of uptake of transferrin-Alexa647 was used as a functional test of knockdown efficacy. (D) Quantification of chemically-induced endocytosis. Fraction of puncta that were green and red is shown for each cell (spots), bars indicate the mean ± 1 s.d. N_exp_ = 3. ANOVA, p = 1.48 × 10^−8^. (E) Western blot to assess depletion of μ2 by RNAi. Cells were prepared in parallel with the experiment shown in C. Blotting for μ2 or for GAPDH (as a loading control) is shown. Scale bars, 10 μm.

### Limited processing of hot-wired endocytic vesicles

What downstream processing occurs to the vesicles that are created by hot-wired CME? To answer this question, we used live imaging of cells co-expressing our system with a variety of organellar markers. Hot-wired CME was triggered by rapamycin application and the colocalization of the markers with bright FKBP-β2-mRuby2 puncta was assessed. No colocalization was seen with markers of early endosomes or late endosomes/lysosomes (Figure 6A). As described above, the puncta colocalize with clathrin, suggesting that the vesicles formed by hot-wiring remain coated with clathrin and do not fuse with early endosomes. In support of this conclusion, we could observe bright FKBP-β2-mRuby2 vesicles “dancing” around a GFP-EEA1 structure for several minutes, but no fusion could be observed (Figure 6B). Perhaps this is due to a lack of SNAREs being packaged into the synthetic vesicles. The limited downstream processing of the hot-wired vesicles suggests that the main use for our system is in studying the stages of endocytosis from initiation to scission but not onward trafficking.

**Figure 6.**
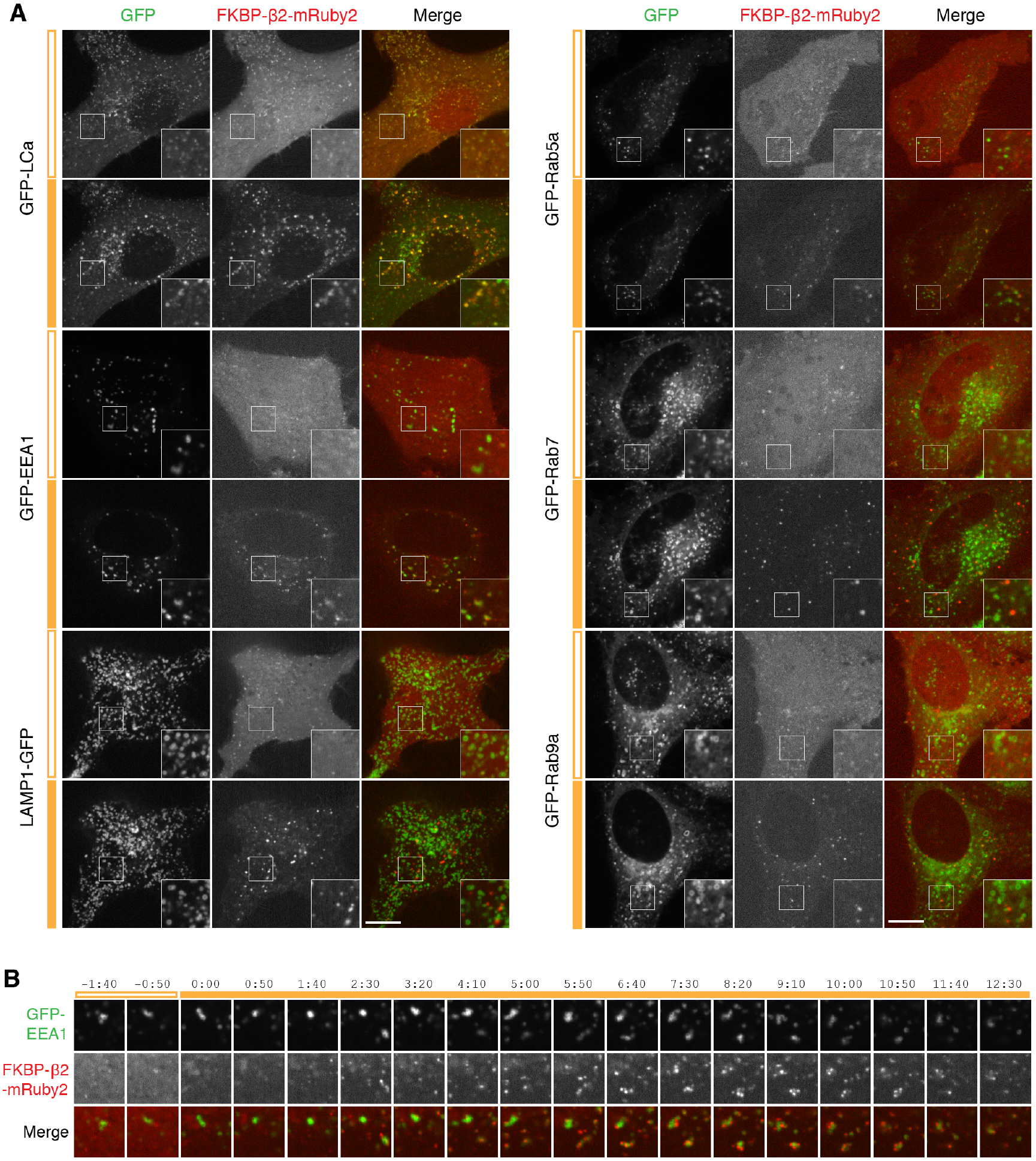
Further trafficking of hot-wired endocytic vesicles is limited. (A) Stills from typical live cell confocal imaging experiments. HeLa cells expressing CD8-dCherry-FRB, FKBP-β2-mRuby2 and the indicated GFP-tagged construct. Endocytosis was triggered by rapamycin addition (200 nM, filled orange bar). For each, upper image shows the frame before rerouting occurs and lower image is 133 frames (665 s) later. Insets show 2X zoom of the boxed area. Scale bar, 10 μm. (B) Gallery of images to show the spatial relationship between chemically-induced endocytic vesicles (red, FKBP-β2-mRuby2) and GFP-EEA1-positive vesicles (green).

### Hot-wired endocytosis cannot override endocytic shutdown mechanisms

One characteristic of CME is that it is inhibited during the early stages of mitosis (Fielding et al., 2012; Kaur et al., 2014). We therefore tested if hot-wired endocytosis could override this inhibition. HeLa cells expressing our system were treated with rapamycin at metaphase. FKBP-β2-GFP relocated to the plasma membrane as usual, however bright green puncta did not form (Figure 7, Supplementary Video S4). Around 30 min later, after the cell had undergone cytokinesis, we began to see puncta formation, showing that induction of endocytosis was ultimately possible, just not during mitosis. We imaged 17 cells in 4 independent experiments. Inhibition of vesicle formation at metaphase and anaphase was observed in 17/17 cells. Of these, 7 cells could be tracked through to cytokinesis, where all 7 began puncta formation. These observations suggest that hot-wiring cannot overcome the mitotic inhibition of CME.

**Figure 7.**
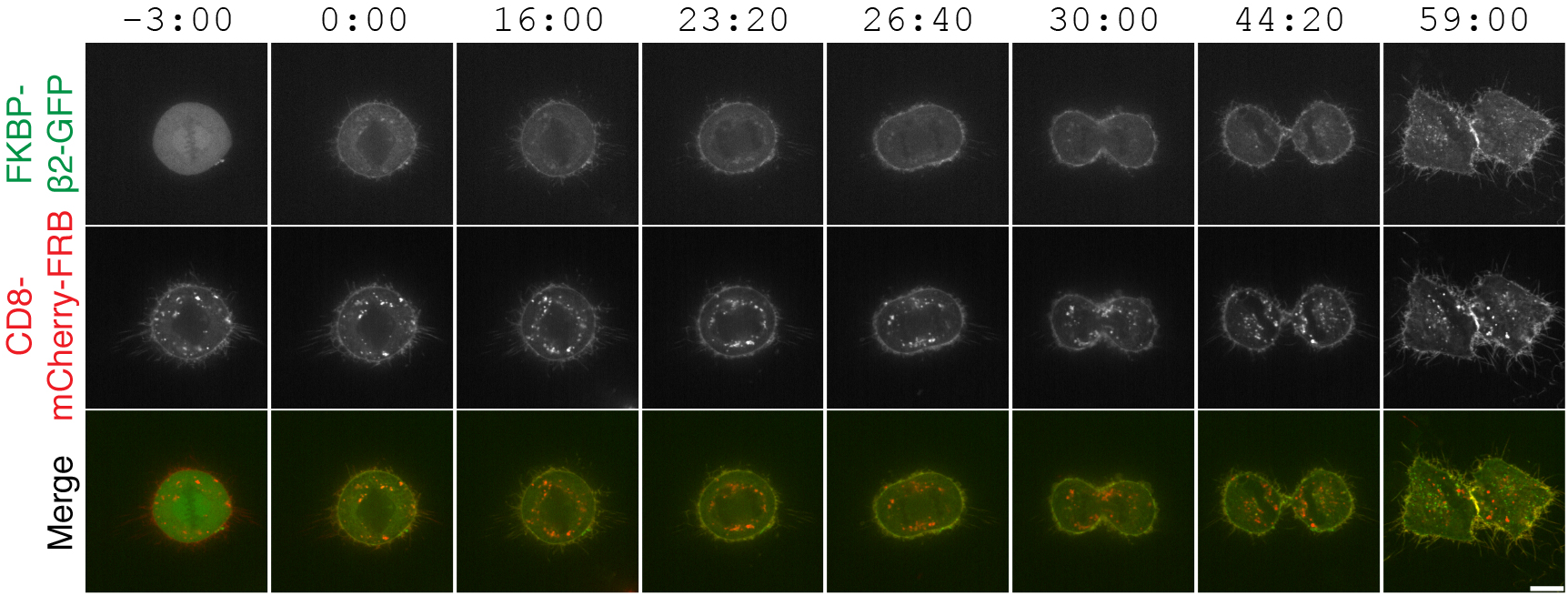
Chemically-induced endocytosis is inhibited during mitosis. Live cell imaging experiment, mitotic cell expressing CD8-mCherry-FRB with FKBP-β2-GFP. Rapamycin (200 nM) was applied during metaphase. Clear rerouting occurs, but no puncta form in the cytoplasm until after cytokinesis. This cell is shown in Supplementary Movie S3. Scale bars, 10 μm.

### Comparison of different clathrin hooks for chemically inducible endocytosis

The hinge and appendage of the β2 subunit of AP2 was initially chosen as our clathrin hook because it is AP2 that coordinates endocytosis at the plasma membrane, however, we hypothesized that the clathrin-binding region from any adaptor would be sufficient to initiate endocytosis. We next compared various clathrin hooks for their ability to initiate endocytosis. Using the same chemically-inducible system, we observed differences in bright green spot formation despite all proteins being efficiently recruited to CD8-mCherry-FRB. Efficient spot formation was seen with the hinge and appendage from β1 subunit of the AP1 complex and with a fragment from epsin which contains clathrin-binding motifs (Figure 8). No spot formation was observed with the hinge and appendage from the α subunit of AP2, which was expected since it does not directly bind clathrin. Surprisingly, the hinge and appendage from the β3 subunit of AP3, which can bind clathrin *in vitro* (Dell’Angelica et al., 1998), was unable to form bright green puncta. Quantification of this behavior showed that β2 was the most efficient and potent initiator, above β1 and epsin, while β3 initiates only marginally more than α adaptin and GFP negative controls (Figure 8B,C). These experiments indicate that one use for this system is to test proteins for *functional* clathrin-binding. Our optimized system for chemically hot-wired endocytosis is therefore a CD8-mCherry-FRB plasma membrane anchor and an FKBP-β2-GFP clathrin hook.

**Figure 8.**
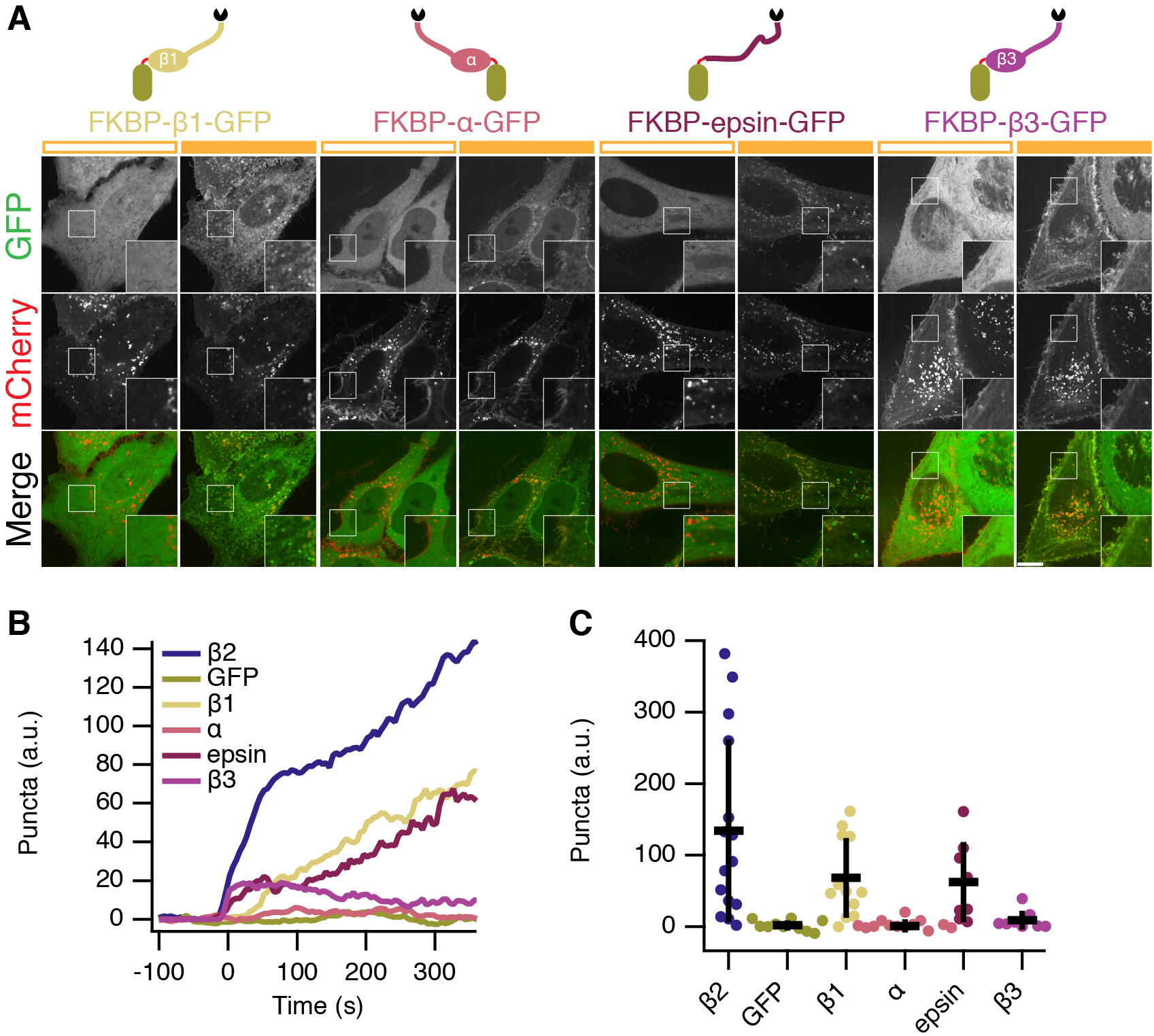
Chemically-inducible internalization: comparison of clathrin hooks. (A) Comparison of four potential clathrin hooks for chemically-inducible endocytosis. Cells expressed CD8-mCherry-FRB with either FKBP-β1-GFP, FKBP-α-GFP, FKBP-epsin-GFP or FKBP-β3-GFP. Stills from live cell imaging experiments are shown, left image shows the frame before rerouting occurs and right image is 133 frames (665 s) later. Inset shows a 2X zoom of the boxed region. Scale bar, 10 m^i. (B) Average plots to show chemically-inducible endocytosis. Colored traces indicate the total area of bright GFP puncta that form in cells after rerouting. Traces were time aligned and averaged. (C) Scatter plot to indicate variability in chemically-induced endocytosis. The total normalized area above threshold at 5 min after rerouting for each cell analyzed is shown (spots). ANOVA, p = 1.142 × 10^−5^. Bars indicate the mean ± 1 s.d. N_cell_=8–15, N_exp_=3.

### Functional CME requires two clathrin interaction sites on AP2β2

We next applied our synthetic system to answer a cell biological question: how does clathrin *functionally* engage with AP2? Two sites on β2 have been implicated in binding clathrin, a clathrin box motif (LLNLD) in the hinge region (Shih et al., 1995), and a tyrosine residue (Y815) in the appendage (Edeling et al., 2006) (Figure 9A). However, it is not clear which sites are important in a functional context. We first confirmed that these two sites interact with clathrin biochemically (Figure 9B). Deletion of the CBM interfered with binding to a CHC fragment containing the terminal domain and ankle region (residues 1–1074). Mutation of Y815A (Y-A) also inhibited binding but to a lesser extent.

**Figure 9.**
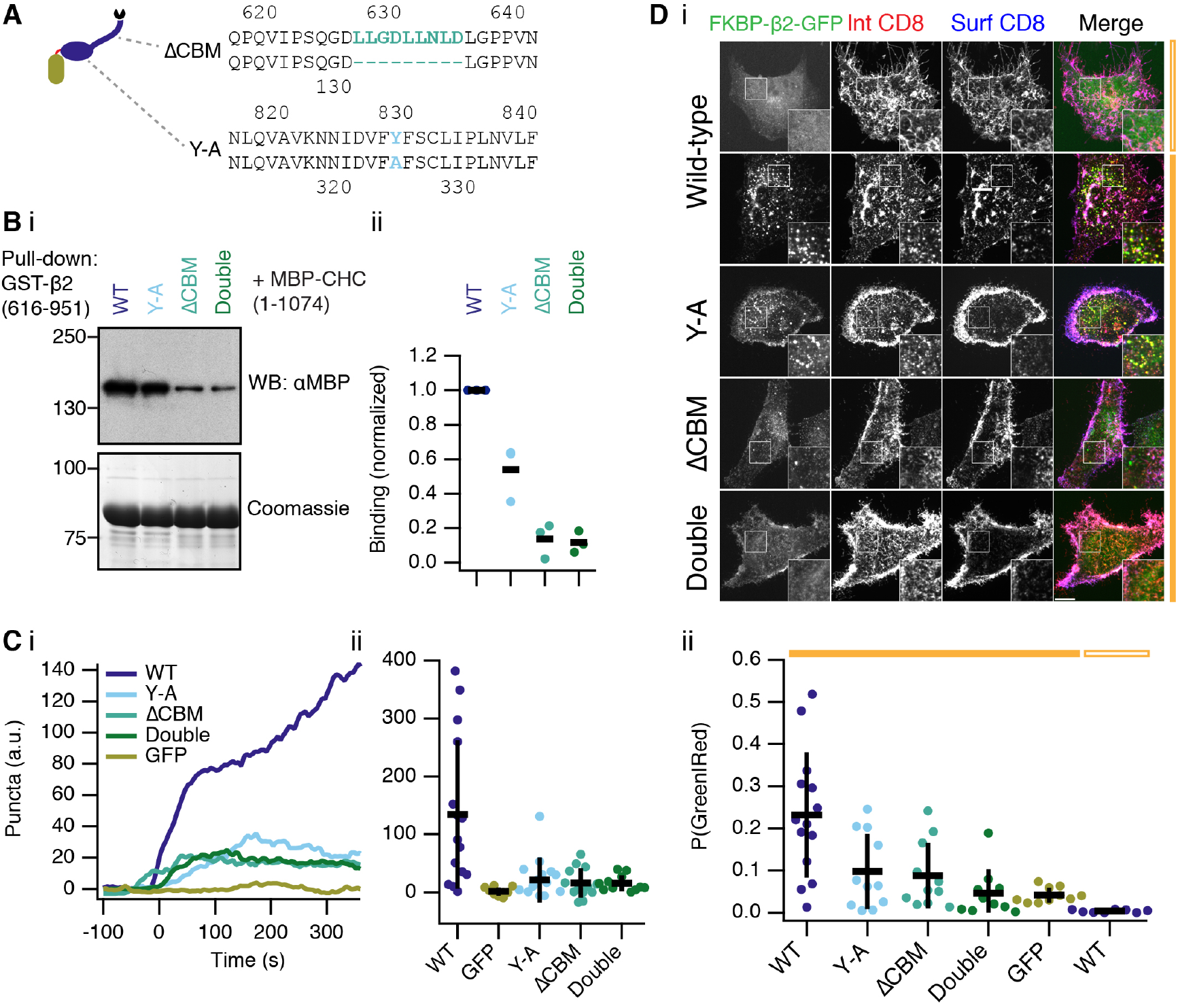
Clathrin functionally engages AP-2 at the plasma membrane using two distinct interactions. (A) Schematic illustration of the putative interactions between β2 and clathrin. Inset shows mutations to delete the clathrin box motif (ΔCBM) or mutation of tyrosine residue Y815 in the appendage (Y-A). Note that Y815 in earlier papers refers to Y829 in our longer isoform. (B) Analysis of clathrin-β2 interaction *in vitro.* Binding experiments using GST-β2(616–951) wild-type or mutants to pull-down MBP-CHC(1–1074) as indicated. Interaction was assessed by western blotting (i) and a coomassie-stained gel was run in parallel to check for equivalent capture on beads. Three independent experiments were performed and analyzed by densitometry (ii). Bar indicates the mean. (C) Quantification of live-cell imaging experiments to show extent of endocytosis triggered by FKBP-β2-GFP or related mutants (see inset). Colored traces indicate the total area of bright GFP puncta that form in cells after rerouting. Traces were time aligned and averaged. Average traces are shown (i) and a scatter plot (ii) of values at 5 min after rerouting for each cell analyzed is shown (spots). Bars indicate the mean ± 1 s.d. N_cell_=11–15, N_exp_=3. ANOVA, p = 7.92 × 10^−5^. Note, the values for FKBP-β2-GFP (WT) and GFP-FKBP (GFP) are also shown in Figure 8. Example live cell movies are shown in Supplementary Figures S5, S6 and S7. (D) Representative images (i) of an immunolabelling experiment to assess endocytosis of CD8 by triggered endocytosis. Scatter plot (ii) to indicate variability in chemically-induced endocytosis. The fraction of green puncta that were also red for each cell analyzed is shown (spots). Bars indicate the mean ± 1 s.d. N_cell_=8–14, N_exp_=3. ANOVA, p = 1.66 × 10^−6^.

Live cell imaging revealed that deletion of the clathrin box motif (ΔCBM) in the hinge of FKBP-β2-GFP and/or mutation of Y815 in the appendage (Y-A) were sufficient to block puncta formation by hot-wiring (Figure 9C, Supplementary Movies S5, S6 and S7). This suggested that both sites were important for internalization. To confirm this result, we used live-cell antibody labeling to further quantify the inhibition (Figure 9D). Internalization of antibody upon rapamycin application was clearly observed in cells expressing wild-type FKBP-β2-GFP with CD8-dCherry-FRB. The degree of internalization was inhibited to a similar extent in cells expressing Y-A or ΔCBM forms of FKBP-β2-GFP. Further inhibition was seen in cells expressing a double mutant FKBP-β2-GFP, such that it was indistinguishable from GFP-FKBP (Figure 9D). These data indicate that both clathrin interaction sites on β2 are required for engagement in a functional context.

### Optogenetic activation of endocytosis

Having successfully implemented temporal control of endocytic initiation using a chemogenetic approach, we next wanted to add spatial control. To do this we used an optogenetic strategy: utilizing the ability of photosensitive LOV2 domain from *Avena sativa* phototropin 1, to cage a small peptide which can bind the engineered PDZ domain (ePDZb1) after exposure to <500 nm light (Strickland et al., 2012; van Bergeijk et al., 2015; Wagner and Glotzer, 2016). We designed light-activated versions of our preferred plasma membrane anchor, CD8-TagRFP657-LOVpep(T406A,T407A,I532A), and clathrin hook, ePDZb1-β2-mCherry, which would allow for discrete dimerization by blue light (Figure 10A). Recruitment of ePDZb1 to the plasma membrane anchor is rapid and reversible (τ = ~40 s, Figure 10B). Transient recruitment was not sufficient to make vesicles. However sustained, patterned illumination of 488 nm light was sufficient to reroute ePDZb1-β2-mCherry to the plasma membrane and to subsequently initiate vesicle formation (Figure 10C). Interestingly once formed, the vesicles did not travel far outside the area of photoactivation (Figure 10D, Supplementary Movie S8). Analysis of optogenetic hot-wiring showed that onset of vesicle formation was fast and was less variable than with the chemical method (Figure 10E). We found ePDZb1 and ePDZb variants of the clathrin hook to be equivalent, and that the LOVpep(T406A,T407A,I532A) version of the plasma membrane anchor was superior to the wild-type. These data show that spatiotemporal control over initiation of endocytosis is possible and is an efficient method to probe questions related to “on demand” traffic from the membrane.

**Figure 10.**
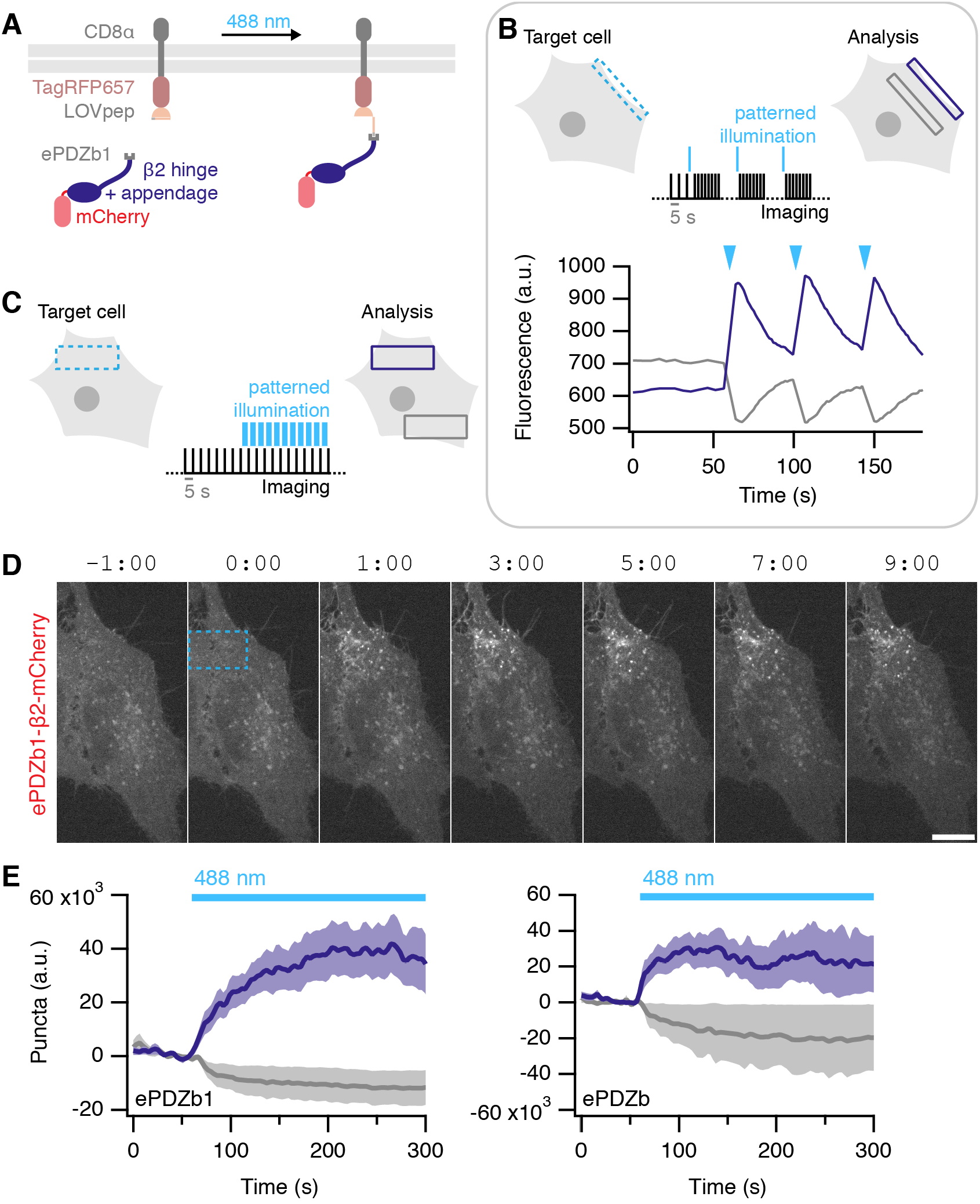
Optically-inducible endocytosis. (A) Illustration of optically-induced endocytosis. Cells co-express a plasma membrane anchored LOVpep (CD8-TagRFP657-LOVpep) and a clathrin hook with a PDZb1 tag (PDZb1-β2-mCherry). Upon illumination with blue light, an epitope is exposed to which the PDZ domain can bind, and the clathrin hook is rerouted to the plasma membrane so that endocytosis can occur at that site. (B) Rapid and reversible ePDZb1-mCherry recruitment to CD8-TagRFP657-LOVpep(T406A,T407A,I532A) at the plasma membrane. Brief (<1 s) patterned illumination of plasma membrane areas followed by rapid imaging. Mean pixel density of an ROI at the membrane (purple) compared to one in the cytoplasm (gray) are shown. (C) Endocytosis is activated in a spatiotemporal manner by sustained, patterned light activation within a defined ROI (blue dashed box). For analysis, the number of vesicles formed within the ROI (purple box) is compared with a similar ROI outside the activated region (gray box). (D) Stills from a typical optogenetic hot-wiring experiment. HeLa cell expressing CD8-TagRFP657-LOVpep(T406A,T407A,I532A) and ePDZb1-β2-mCherry. Only the mCherry channel is shown. Time 0 indicates when patterned illumination began. A movie of this panel is available as Supplementary Movie S8. (E) Plots of mean ± s.e.m. vesicle formation in cells expressing CD8-TagRFP657-LOVpep(T406A,T407A,I532A) and either ePDZb1-β2-mCherry (n_cell_ = 13, n_exp_ = 4) or ePDZb-β2-mCherry (n_cell_ = 8, n_exp_ = 3) as indicated. Purple and gray indicate vesicles formed inside and outside the photo-activation zone, respectively.

## Discussion

In this paper we described systems for triggering endocytosis on demand. These tools are useful because they allow the internalization of defined cargo with temporal or spatiotemporal control via chemical or optical methodology, respectively. Bypassing several regulatory steps in constitutive CME to hot-wire endocytic events has many potential uses and here we used it to demonstrate how clathrin and AP2 interact functionally to form a CCV and to test the mitotic inhibition of CME.

Clathrin-coated pits are initiated when AP2 that is bound to PI(4,5)P_2_ at the plasma membrane simultaneously binds cargo bearing AP2-interacting motifs (Traub, 2009). This exposes the β2 hinge and appendage in order to bind clathrin and begin polymerization and stabilization of the nascent CCP (Kelly et al., 2014; Kirchhausen et al., 2014). Vesicle formation then proceeds in a regulated manner (Aguet et al., 2013; Loerke et al., 2009; Mettlen et al., 2009; Taylor et al., 2011). Our synthetic systems hot-wire this process by directly recruiting a clathrin hook to the plasma membrane. This mimicking of the PI(4,5)P_2_-bound, cargo-stabilized form of AP2, eliminates all the preceding steps, and immediately starts clathrin polymerization. Hot-wired CME is unlikely to work in isolation from the rest of the endocytic machinery. Most obviously, other components such as dynamin are required to form a free hot-wired CCV. Moreover, hot-wired CME is still under the same regulation as endogenous CME. We found that hot-wiring cannot overcome the mitotic inhibition of CME. Interestingly, hot-wired events only occurred after mitosis completion, whereas endogenous CME restarts in late anaphase (Fielding and Royle, 2013; Schweitzer et al., 2005). This suggests that hot-wired CME may be more sensitive to changes in membrane tension and actin availability that govern the mitotic inhibition of CME (Kaur et al., 2014).

Since hot-wired events must share resources with endogenous CME, can hot-wiring be considered orthogonal to it? We found no profound impact of hot-wiring on endogenous CME. Our conclusion is that there are sufficient resources in the cell to support endogenous CME as well as the additional, triggered form of CME. This is perhaps to be expected since cells often have to increase the rate of CCP formation when certain receptors are activated (see Reeves et al., 2016 for a recent example), and this is analogous to hot-wiring. There may be more subtle changes to endogenous CME as a result of hot-wiring. Precisely how, and to what extent, hot-wiring interfaces with endogenous CME needs further examination. We note however, that potential interference with endogenous CME is less of a concern with optogenetic hot-wiring, because this system is rapidly reversible.

When testing orthogonality of our system, we found evidence that hot-wired CME can occur independently of endogenous AP2. However, we cannot conclude that AP2 is dispensable for hot-wired CME. First, it is notoriously difficult to eliminate AP2 by RNAi (Boucrot et al., 2010). Although our depletion was sufficient to block transferrin uptake, it is possible that trace levels of endogenous AP2 still help the hot-wired vesicles to form. Second, we think it is most likely that AP2 is involved after hot-wiring has occurred since AP2 has a clear regulatory role in coordinating CME (Aguet et al., 2013). In any case, the likely application of our systems will be in normal cells with a full complement of AP2 and other endocytic proteins. So, while it might be possible to trigger endocytosis without AP2, this is not relevant to the general application of our system.

One application of hot-wiring endocytosis is to test for *functional* interactions between clathrin and clathrin-binding proteins. It’s clear that binding between protein fragments in a test tube does not necessarily translate to functional interactions in cells. Even *in situ* we have previously seen that colocalization of adaptors and clathrin mutants does not reliably indicate functional CME (Willox and Royle, 2012). The inducibility of hot-wiring allows the assessment of functional interactions in ways that expression of mutants do not. We used this technique to demonstrate that two necessary interactions between AP2 and clathrin are both required for CME. A clathrin box motif in the β2 hinge binds to the terminal domain, and a site on the β2 appendage interacts with a distinct site probably on the ankle of clathrin heavy chain. Biochemical evidence for these interactions was in the literature (Edeling et al., 2006; Hood et al., 2013; Owen et al., 2000; ter Haar et al., 2000), but the relative importance of each site for function was previously unknown. A further surprising result was that β3, which is known to bind clathrin, was not interchangeable with β1 and β2 as a functional initiator of CME. The β3-clathrin interaction is robust since clathrin box motifs were actually discovered using β3 (Dell’Angelica et al., 1998). Subsequent work questioned whether AP3 was a genuine clathrin adaptor (Peden et al., 2002; Zlatic et al., 2013). Our results suggest that functional clathrin engagement by β3 is not equivalent to β1 and β2. One possible explanation is that clathrin hooks must also interact with endocytic accessory proteins for proper coat maturation and that β3 is unable to do this (Schmid et al., 2006).

While our systems are ready-to-use, we think further optimization of the plasma membrane anchor may be possible. First, some internal vesicles, which contained membrane anchor, were present with CD8-, CD4-and GAP43-based anchors. These internal vesicles complicate image analysis and might prevent efficient rerouting exclusively to the plasma membrane. Second, anchors are prone to diffusion in the plasma membrane. For the optogenetic application, this means that we constantly lose photoactivated anchors. Although the anchors at the plasma membrane quickly inactivate once they diffuse out of the illumination zone which ensures local activation, this behavior reduces the efficiency of vesicle creation. Engineering the anchors to address these issues are key to further improvements in the technology. Moreover, diversifying the cargoes that are programmed for internalization will further extend the capabilities of hot-wiring.

There are numerous uses for our hot-wiring systems. For example, the possible functionalization of the plasma membrane anchor on the extracellular face so that targeted uptake of nanoparticles or other interesting materials is possible. Another short-term goal of ours is to change cell behavior by local optogenetic activation of endocytosis. In addition, we anticipate that these systems will be useful for understanding initiation of endocytic events as well as downstream processing of hot-wired vesicles, and the machineries that control these aspects of vesicle trafficking.

## Methods

### Cell culture

HeLa cells (HPA/ECACC #93021013) were maintained in Dulbecco’s Modified Eagle’s Medium (DMEM) plus 10% fetal bovine serum (FBS) and 100 U/ml penicillin/streptomycin. RPE1 cells were maintained in DMEM/F12 supplemented with 10% FBS, 100U/ml penicillin/streptomycin, 0.26% NaHCO3 and 2 mM glutamine. All cells were kept at 37 °C and 5% CO2. DNA transfections were performed with GeneJuice (MerckMillipore, UK) and siRNA transfections with Lipofectamine 2000 (Life Technologies, UK) according to the manufacturer’s instructions. Cells were imaged or fixed 2 days after DNA transfection, siRNA was used in a “two hit” protocol with transfection at 2 and 4 days prior to use (Willox and Royle, 2012). Sequences targeting μ2 (μ2-2) or clathrin heavy chain (chc-2) were as described previously (Motley et al., 2003).

### Molecular biology

CD8-β2-mCherry fusion protein was made by insertion at EcoRI and BamHI of CD8-8A (Fielding et al., 2012) into β2-mCherry, which had been previously prepared from PCR of GST-β2 adaptin(616–951) (Hood et al., 2013) inserted into pmCherry-N1. The non-fluorescent variant was produced by replacing β2-mCherry with β2 at BspEI and NotI sites. CD8-FRB was made by PCR amplification of CD8-8A and subcloning into pMito-FRB at AgeI and EcoRI; CD8-mCherry-FRB was made from CD8-FRB by replacing FRB with mCherry-FRB from pMito-mCherry-FRB at the same sites. CD4-mCherry-FRB was made by overlap extension PCR of CD8-mCherry-FRB and CD4-FRB (synthesized by IDT); the third membrane anchor, GAP43-FRB-RFP, was available from previous work. Dark versions of mCherry constructs (dCherry) were made by site-directed mutagenesis (SDM) to introduce the K70N mutation.

FKBP-β2-mCherry was made by inserting FKBP PCR product into β2-mCherry at Acc65I and XhoI sites, this was then modified by subcloning of GFP to make FKBP-β2-GFP. Mutations were added by SDM, Y-A by mutation of β2 adaptin residue Y815 and ΔCBM by deletion of region 627–635 (LLGDLLNLD). The double mutant (ΔCBM/Y-A) was created by switching the β2 adaptin fragment containing the Y815A mutation for the same region in ΔCBM construct using PspOMI and AgeI. Additional FKBP constructs were made by inserting a PCR product in the place of β2 in FKBP-β2-GFP; mouse a adaptin 1 (region 740–977) and mouse epsin (region 144–575) were inserted at Acc65I and AgeI, and human β1 adaptin (region 617–949) was added at AgeI and EcoRI. FKBP-β3-GFP was made by insertion of human β3 fragment (region 702–1094) into FKBP-β1-GFP. FKBP-β2-mRuby2 was made by subcloning of mRuby2 (gift from Joachim Goedhart, Amsterdam) PCR product into FKBP-β2-GFP at AgeI and NotI. GFP-FKBP was available from previous work (Cheeseman et al., 2013).

Endocytic compartment markers were all available from previous work or sourced from Addgene, including mCherry-LCa (Hood and Royle, 2009), GFP-LCa (Royle et al., 2005), GFP-EEA1 (Addgene #42307), and LAMP1-mGFP (Addgene #34831). Dynamin2-GFP was a gift from Richard Vallee (Columbia) GFP-Rab5a, GFP-Rab7, and GFP-Rab9a were gifts from Francis Barr (Oxford).

For optogenetic experiments, pTagRFP657-LOVpep(T406A,T407A,I532A) was made by ligation of LOVpep(T406A,T407A,I532A), ordered as a custom gene from IDT, and pTagRFP657 (Addgene #31959). This product was then inserted into CD8-mCherry at EcoRI and AgeI sites. PDZb-β2-GFP and PDZb1-β2-GFP were made by PCR of PDZb (Addgene #34980) or PDZb1 (Addgene #34981), inserted into FKBP-β2-GFP at XhoI and BamHI, replacing FKBP. PDZb-GFP and PDZb1-GFP were made by subcloning of PDZb or PDZb1 (different PCR products to preserve reading frame) into pEGFP-N1. PDZb-β2-mCherry and PDZb1-β2-mCherry were made by subcloning of PDZb/PDZb1 from PDZb-β2-GFP or PDZb1-β2-GFP and inserting into FKBP-β2-mCherry at BamHI and XhoI sites. PDZb-mCherry and PDZb1-mCherry were made by PCR of PDZb/PDZb1-GFP inserted into pmCherry-N1 at BamHI and AgeI. Plasmids to express MBP-CHC(1–1074)-His6 and GST-β2(616–951) in bacteria were available from previous work (Hood et al., 2013). The GST-β2(616–951) Y-A, ΔCBM and double mutants were made by inserting into pGEX-6P-1 PCR fragments amplified from CD8-β2-mCherry constructs containing these mutations at EcoRI-NotI.

### Immunofluorescence

For live immunolabeling of CD8-β2-mCherry in HeLa cells, 1:100 Alexa488-conjugated mouse anti-CD8 (AbD Serotec) was added at 4 °C or 37 °C for 40 min, diluted in DMEM with 10% FBS. The cells were then placed on ice, washed with PBS with 1% BSA and surface fluorescence quenched with rabbit anti-Alexa488 (Invitrogen) before relabeling with 1:500 anti-mouse Alexa633 (Life Technologies) conjugated secondary antibody.

Live labeling of rapamycin-inducible system was performed in HeLa cells transfected with CD8-mCherry-FRB and FKBP-β2-GFP (or mutant variants) or FKBP-GFP. Live cells were incubated with untagged anti-CD8 (AbD Serotec) at 1:1000 at 37 °C for 40 min; 200 nM rapamycin was then added for 30 min before cells were put on ice and washed with PBS with 1% BSA. Alexa647-conjugated anti-mouse secondary was added in DMEM at 1:500 for 1 h then cells were fixed and permeabilized with 0.1% Triton X-100 in PBS. Finally, 1:500 Alexa568-conjugated secondary antibody was added in DMEM for 1 hour before washing in PBS and mounting.

Live cell imaging of antibody uptake was performed in cells transfected with FKBP-β2-GFP or GFP-FKBP with CD8-mCherry-FRB or CD8-dCherry-FRB + mCherry LCa. Alexa647-conjugated anti-CD8 (AbD Serotec) was added to HeLa cells at 37 °C for 30 min and cells washed twice in warm medium before imaging using the standard live-cell imaging protocol described below.

For transferrin uptake analysis, HeLa cells were serum-starved for 20 min in serum-free DMEM then exposed to 200 nM rapamycin or ethanol vehicle for 20 min, with Alexa647-conjugated transferrin (Invitrogen) added for the final 10 min before fixing. All dilutions were performed in serum-free media.

In experiments where both transferrin uptake and live antibody labeling were performed, cells were first starved for 30 min in serum-free DMEM. Mouse anti-CD8 was applied for 40 min in serum-free DMEM, then cells were treated with rapamycin (200 nM) or vehicle for 30 min. Alexa647-conjugated transferrin was applied for the last 10 min of treatment before cells were moved to ice. Unconjugated goat anti-mouse antibody was added to quench surface anti-CD8. Cells were then fixed with PFA, permeabilized and then incubated with Alexa568-conjugated secondary antibody in DMEM for 1 hour before washing in PBS and mounting.

### Light microscopy

For live-cell imaging of rerouting experiments, HeLa cells were transfected with CD8-mCherry-FRB, CD8-dCherry-FRB, CD4-mCherry-FRB, pMito-mCherry-FRB or GAP43-FRB-mRFP with FKBP-β2-GFP (or mutants), FKBP-β2-mRuby2, FKBP-β1-GFP, FKBP-α-GFP, FKBP-β3-GFP, FKBP-epsin-GFP or GFP-FKBP. For experiments where the red channel was needed for imaging something other than CD8, dark variants of mCherry constructs were used where mCherry had been mutated to give K70N (Subach et al., 2009), we refer to this variant as dCherry (dark mCherry). Cells were imaged in glass-bottom fluorodishes (WPI) with Leibovitz L-15 CO_2_-independent medium (Sigma) supplemented with 10% FBS and kept at 37 °C. Imaging was performed using a spinning disc confocal system (Ultraview Vox, Perkin Elmer) with a 100x 1.4 NA oil-immersion objective. Images were captured every 5, 20 or 30 s using a dual camera system (Hamamatsu ORCA-R2) after excitation with lasers of wavelength 488 nm, 561 nm, and 640 nm (if required); 200 nM rapamycin (Alfa Aesar) was added after 1 or 2 min.

Light-induced dimerization experiments were performed similarly, cells transfected with CD8-TagRFP657-LOVpep(T406A,T407A,I532A) and PDZb1-β2-mCherry or PDZb-β2-mCherry were imaged with 561 nm and 640 nm lasers, captured with a single camera at 5 s intervals. Photoexcitation was done using a 488 nm laser in a defined ROI. 12 frames of baseline were captured before interleaving photoactivation and imaging, as indicated in Figure 8C. Use of the CA form of LOVpep generated many vesicles similar to our CD8-β2-mCherry fusion protein.

All fixed cell and immunostaining experiments were performed in transiently transfected HeLa cells fixed in PBS with 3% paraformaldehyde, 4% sucrose and mounted in Mowiol with DAPI. Imaging was performed using the same Ultraview spinning disc confocal and 100x objective, z-slices were taken at 0.5 miti intervals.

TIRF microscopy was performed on RPE1 cells expressing CD8-dCherry-FRB and FKBP-β2-mRuby2 with GFP-LCa or Dynamin2-GFP. Images were captured at 1 frame/s using an Olympus IX81 equipped with a 100x 1.49 NA objective and Hamamatsu ImagEM-1K EMCCD following excitation with 488 nm and 561 nm lasers. 200 nM rapamycin was added after 1 min.

### Correlative light-electron microscopy (CLEM)

HeLa cells transfected with CD8-dCherry-FRB and FKBP-β2-GFP or GFP-FKBP were imaged in gridded glass bottom dishes (P35G-2-14-CGRD, MatTek Corp, Ashland, MA) following incubation with anti-CD8 (1:1000) for 30 min and then Alexa-546 FluoroNanoGold™-anti-mouse Fab’ (1:200) for 10 min. Using the photo-etched coordinates on each grid, the cell location was recorded using brightfield illumination at 20x. Following Rapamycin addition, fluorescent live cell imaging determined vesicle formation at 100x magnification on a Nikon Ti epifluorescence microscope and a Coolsnap Myo camera (Photometrics) using NIS Elements AR software. Once a sufficient number of vesicles had been observed, the sample was immediately fixed with 3% glutaraldehyde and 0.5% paraformaldehyde in 0.05M phosphate buffer pH 7.4 for 2 hours. Remaining aldehydes were quenched in 50 mM glycine and cells were subsequently washed in dH_2_O. Gold enhancement was then performed for 3 min as per manufacturer’s instructions (GoldEnhance-EM, Nanoprobes, Inc.). Cells were washed twice in 0.3% Na_2_S_2_O_3_ and then thoroughly in dH_2_O. Gold-enhanced samples were postfixed in 1% OsO_4_ for 60 min, rinsed with dH_2_O and then stained *en bloc* with 0.5% uranyl acetate in 30% ethanol for 60 min. Cells were dehydrated in increasing concentrations of ethanol for 10 min each (50%, 70%, 80%, 90%, 100%, 100%) before being embedded in epoxy resin (TAAB) and left to polymerize for 48 h at 60 °C. Each cell of interest was identified by correlating the grid and cell pattern on the surface of the polymerized block with previously acquired brightfield images. Ultrathin (70 nm) sections were collected on formvar coated copper grids using an EM-UC6 ultra-microtome (Leica Microsystems) and contrasted with saturated aqueous uranyl acetate and Reynolds lead citrate. Sections were imaged on a Jeol 1400 TEM at 100 kV.

### Biochemistry

Binding experiments to test the interaction between β2 and clathrin were carried out as previously described (Hood et al., 2013). Briefly, GST-β2 adaptin(616–951)-His6 and MBP-CHC(1-1074)-His6 proteins were purified using glutathione sepharose 4B and amylose resin, respectively. For *in vitro* interaction studies, equal amounts (50 μg) of GST- and MBP-fused proteins were mixed in reaction buffer I (50 mM Tris-Cl pH 7.5, 150 mM NaCl, 0.1 mM EGTA). The mixture was incubated with a 50% slurry of glutathione sepharose 4B beads (pre-equilibrated in NET-2 buffer ‘50 mM Tris-Cl pH 7.5, 150 mM NaCl, 0.5% NP-40 substitute]) and left overnight at 4°C with rotation. Next day, beads were collected by spinning down at 1,000 × *g* for 2 min at 4°C and washed 4 times with NET-2 buffer. Beads were then resuspended in 30 μl of 1X Laemmli buffer, denatured at 95°C and analyzed by western blotting after running on 8% SDS-PAGE. Protein samples were also analyzed by staining SDS-PAGE gels with coomassie brilliant blue. Antibodies used for western blotting: anti^2 (mouse anti-AP50 Clone 31, BD Transduction Labs), anti-GAPDH (rabbit G9545, Sigma), anti-MBP (mouse 8G1 2396; Cell Signaling Technology).

### Image analysis

For analysis of triggered endocytosis, a simple thresholding method proved the most straightforward. The green channel was first corrected for photobleaching using the simple ratio method. To examine rerouting kinetics, the mean pixel density within a cytoplasmic ROI was measured for all frames in the movie. Next, a binary threshold was applied to the green channel which isolated any newly formed vesicles and excluded background signal from other structures such as the plasma membrane. The raw integrated density (number of pixels over threshold * 255) within an ROI containing the cell were measured for all frames in the movie. These values for all cells were fed into Igor Pro 7 and a series of custom-written functions processed the data (available at https://github.com/quantixed/PaperCode/). The raw integrated densities were aligned to time 0, which was defined as the point at which the mean pixel density of the cytoplasmic ROI fell by 50%. These time-aligned traces were normalized by the size of the ROI and baseline subtracted. This gave a good approximation of the number of internalized vesicles as defined by bright puncta of GFP-clathrin hook-FKBP fluorescence. For light-activated endocytosis the same method was used with no time alignment. The raw integrated density was extracted in the same way except measurements were taken within the illuminated ROI and an equivalent, non-exposed area of the same cell. Due to variability in image capture, time-stamps of image capture were extracted from the mvd2 library using IgorPro. These were used for interpolation to calculate precise averages in IgorPro.

For analysis of antibody feeding experiments, four z slices from the centre of the stack were thresholded in the red and green channels to isolate vesicular structures. Using Fiji and the ’analyze particles’ plugin, a mask showing only particles of 0.03–0.8 m! and circularity of 0.3–1.0 was created. Particles present in both channels were determined using the image calculator and a third mask created. The pixels over threshold within each processed cell mask were measured for red+green and for red alone, and the values combined from each of the z slices. The ratio of red+green/red pixels was the proportion of red puncta which were GFP-positive, discarding structures containing only CD8. Transferrin uptake was measured using the same method and parameters, using results from the transferrin channel alone.

Analysis of TIRFM images was performed in MATLAB using the cmeAnalysis package (Aguet et al., 2013). Recruitment of GFP-dynamin2 or LCa-GFP to FKBP-β2-mRuby2 positive puncta was observed using the red channel as the master and green as the slave. Additionally, kymographs were created in Fiji.

Figures were made in Fiji, Igor Pro, Photoshop and assembled in Illustrator. Statistical tests between two groups was done using Student’s t-test with Welch’s correction; between three or more groups, one-way ANOVA was used with Tukey’s post-hoc test.

## Acknowledgements

We would like to thank Zuzana Kadlecova for extensive discussions and experimental work not described here, and Sandra Schmid for constructive comments on our manuscript. We thank: Anne Straube for access to TIRF microscopy; Rebecca Milton, Rachel Jones and Aditi Kibe for technical assistance; Joachim Goedhart, Francis Barr and Richard Vallee for reagents. Finally, all Royle Lab members and CMCB colleagues gave valuable comments throughout the project. LAW was supported by a studentship from the Medical Research Council (MR/J003964/1).

## Author contributions

LAW: imaging, analysis, making reagents, and writing the paper

NIC: CLEM imaging SS: binding experiments

SJR: analysis, writing code, and writing the paper.

The authors declare no conflict of interest

---

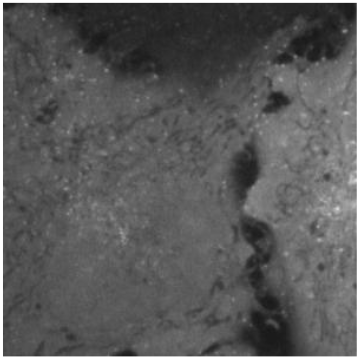
Supplementary Movie S1 Chemical induction of endocytosis: CD8-mCherry-FRB and FKBP-β2-GFP. **Example confocal live cell imaging of rapamycin (200 nM) application to HeLa cells expressing CD8-mCherry-FRB and FKBP**-β**2-GFP. Only the green channel is shown, in grayscale. See Figure 2. Imaging rate, 0.2 fps. Video speed, 10 fps.**

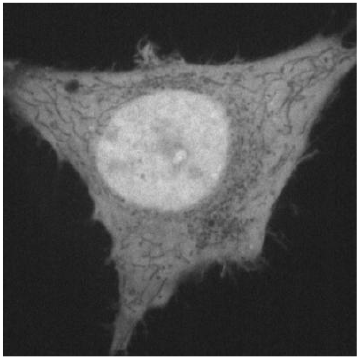
Supplementary Movie S2 No chemical induction of endocytosis: CD8-mCherry-FRB and GFP-FKBP. Example confocal live cell imaging of rapamycin (200 nM) application to HeLa cells expressing CD8-mCherry-FRB and GFP-FKBP. Only the green channel is shown, in grayscale. See Figure 2. Imaging rate, 0.2 fps. Video speed, 10 fps.

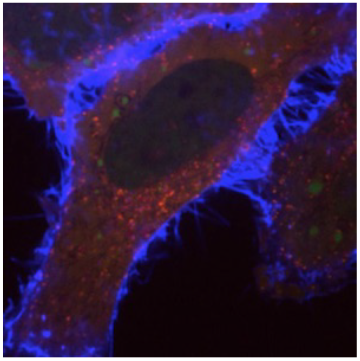
Supplementary Movie S3 Chemical induction of endocytosis: CD8-dCherry-FRB, FKBP-β2-mRuby2 and GFP-LCa with anti-CD8/Alexa647 labeling. **Example confocal live cell imaging of rapamycin (200 nM) application to a mitotic HeLa cell expressing CD8-dCherry-FRB, FKBP**-β**2-mRuby2 (red) and GFP-LCa (green). Anti-CD8/Alexa647 (blue) was added to label the plasma membrane anchor and then rapamycin was applied. See Figure 2. Imaging rate, 0.2 fps. Video speed, 10 fps.**

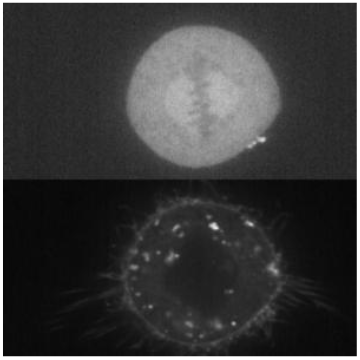
Supplementary Movie S4 Inhibition of hot-wired endocytosis in mitotic cells. Example confocal live cell imaging of rapamycin (200 nM) application to a mitotic HeLa cell expressing CD8-mCherry-FRB and GFP-FKBP. Green channel is above, in grayscale; red channel is shown below. See Figure 7. Imaging rate, 0.05 fps. Video speed, 10 fps.

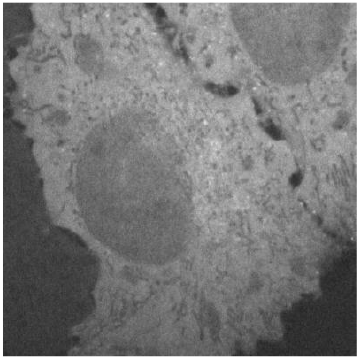
Supplementary Movie S5 Impairment of chemically induced endocytosis using CD8-mCherry-FRB and FKBP-β2-GFP(Y-A) Example confocal live cell imaging of rapamycin (200 nM) application to HeLa cells expressing CD8-mCherry-FRB and FKBP**-**2-GFP(Y-A). Only the green channel is shown, in grayscale. Imaging rate, 0.2 fps. See Figure 9. Video speed, 10 fps.

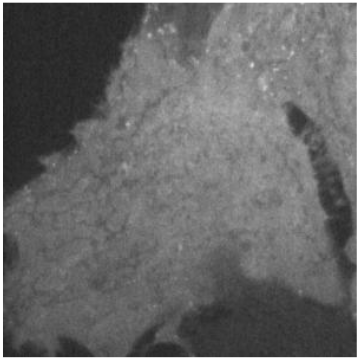
Supplementary Movie S6 Impairment of chemically induced endocytosis using CD8-mCherry-FRB and FKBP-β2-GFPΔCBM) **Example confocal live cell imaging of rapamycin (200 nM) application to HeLa cells expressing CD8-mCherry-FRB and FKBP**-β**2-GFP(ΔCBM). Only the green channel is shown, in grayscale. See Figure 9. Imaging rate, 0.2 fps. Video speed, 10 fps.**

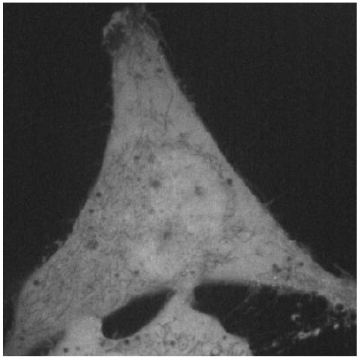
Supplementary Movie S7 Impairment of chemically induced endocytosis using CD8-mCherry-FRB and FKBP-β2-GFP(double mutant) Example confocal live cell imaging of rapamycin (200 nM) application to HeLa cells expressing CD8-mCherry-FRB and FKBP**-β**2-GFP(double mutant). Only the green channel is shown, in grayscale. See Figure 9. Imaging rate, 0.2 fps. Video speed, 10 fps.?

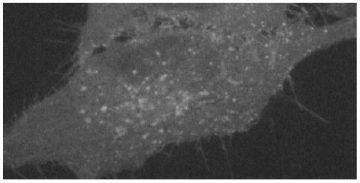
Supplementary Movie S8 Optical induction of endocytosis: CD8-TagRFP657-LOVpep(T406A,T407A,I532A) and PDZb1-β2-mCherry. Example confocal live cell imaging of light (488 nm) activation of HeLa cells expressing TagRFP657-LOVpep(T406A,T407A,I532A) and PDZb1-β2-mCherry. Only the mCherry channel is shown, in grayscale. Patterned illumination was done as described in Figure 10. Imaging rate, 0.2 fps. Video speed, 10 fps.

